# Identification of a rapidly-spreading triple mutant for high-level metabolic insecticide resistance in *Anopheles gambiae* provides a real-time molecular diagnostic for anti-malarial intervention deployment

**DOI:** 10.1101/2021.02.11.429702

**Authors:** Harun Njoroge, Arjen van’t Hof, Ambrose Oruni, Dimitra Pipini, Sanjay C. Nagi, Amy Lynd, Eric R. Lucas, Sean Tomlinson, Xavi Grau-Bove, Daniel McDermott, Francis T. Wat’senga, Emile Z. Manzambi, Fiacre R. Agossa, Arlette Mokuba, Seth Irish, Bilali Kabula, Charles Mbogo, Joel Bargul, Mark J.I. Paine, David Weetman, Martin J. Donnelly

## Abstract

Insecticide resistance provides both an increasingly pressing threat to the control of vector-borne diseases and insights into the remarkable capacity of natural populations to show rapid evolutionary responses to contemporary selection. Malaria control remains heavily dependent on deployment of pyrethroid insecticides, primarily in long lasting insecticidal nets (LLINs), but resistance in the major malaria vectors has increased over the last 15 years in concert with dramatic expansion of LLIN distributions. Identifying genetic mechanisms underlying high-level resistance in mosquitoes, which may almost entirely overcome pyrethroid efficacy, is crucial for the development and deployment of potentially resistance-breaking tools. Using the *Anopheles gambiae* 1000 genomes (Ag1000g) data we identified a very recent selective sweep in mosquitoes from Uganda which localized to a cluster of cytochrome P450 genes, including some commonly implicated in resistance. Further interrogation revealed a haplotype involving a trio of mutations, a nonsynonymous point mutation in *Cyp6p4* (I236M), an upstream insertion of a partial Zanzibar-like transposable element (TE) and a duplication of the *Cyp6aa1* gene. The mutations appear to have originated recently in *An. gambiae* from the Kenya-Uganda border region around Lake Victoria, with stepwise replacement of the double-mutant (Zanzibar-like TE and *Cyp6p4-236M*) with the triple-mutant haplotype (including *Cyp6aa1* duplication), which has spread into the Democratic Republic of Congo and Tanzania. The triple-mutant haplotype is strongly associated with increased expression of genes able to metabolise pyrethroids and is strongly predictive of resistance to pyrethroids most notably deltamethrin, a commonly-used LLIN insecticide. Importantly, there was increased mortality in mosquitoes carrying the triple-mutation when exposed to nets co-treated with the synergist piperonyl butoxide (PBO). Frequencies of the triple-mutant haplotype remain spatially variable within countries, suggesting an effective marker system to guide deployment decisions for limited supplies of PBO-pyrethroid co-treated LLINs across African countries. Duplications of the *Cyp6aa1* gene are common in *An. gambiae* across Africa and, given the enzymes metabolic activity, are likely to be a useful diagnostic for high levels of pyrethroid resistance.

## Introduction

Insecticide resistance in disease vectors has become an influential model for understanding rapid contemporary evolution but, more importantly, identifying how resistance arises and spreads is crucial for disease control. Resistance to pyrethroid insecticides in African malaria vector mosquitoes has spread to near ubiquity [1, 2] and, though it is often difficult to demonstrate its impact on malaria infections [3], in some cases it has reached levels that threaten the effectiveness of vector control programmes [4, 5]. A better understanding of resistance distribution and mechanisms will permit a more informed selection and deployment of insecticides to combat evolving mosquito populations. Whilst our understanding of the genetic basis of insecticide resistance in mosquitoes has advanced substantially [6], especially in the important vector *Anopheles funestus* [7], molecular diagnostics for the major vectors in the *An. gambiae* complex remain limited to a handful of mutations[8] which explain a relatively small fraction of the variance in phenotype [9, 10] or which are now at such high frequency as to provide limited diagnostic resolution [11].

Long-lasting insecticidal nets (LLINs) are the principal tool for vector control to combat malaria, especially in sub-Saharan Africa [12]. The majority of LLINs are treated only with pyrethroid insecticides, to which resistance is now widespread [13]. Though behavioural variation and physical or physiological modifications affecting insecticide uptake may sometimes play a role, pyrethroid resistance is caused predominantly by two distinct mechanisms. The first is resistance via point mutations in the target-site of the insecticide, for pyrethroids the Voltage-gated sodium channel (*Vgsc*), which results in decreased sensitivity to the insecticide [14]; the second is metabolic resistance due to over-expression or altered activity of detoxification enzymes, of which the cytochrome P450 family is commonly considered most important [15, 16]. Cytochrome P450 activity is inhibited by the synergist piperonyl-butoxide (PBO), and bed nets incorporating PBO are effective against P450-mediated resistance, as demonstrated by large-scale field trials [4, 17]. Given the continued operational use of pyrethroid-containing nets, it is vital that we understand the genetic mechanisms that may impact their efficacy, to optimise bednet deployment, preferably using information from rapidly-applied DNA markers. In advance of a randomized control trial of PBO-LLINs [4], we sought to characterise pyrethroid resistance mechanisms in the primary malaria vector An. *gambiae s.s*. [18] in Uganda and Kenya.

The recent development of the *Anopheles gambiae* 1000 genomes project (Ag1000g) has led to a step change in our ability to identify DNA variation driven by selection pressure. We have been able to perform genome-wide searches for regions under recent natural selection in insecticide resistant populations across Africa, and work has shown that the strongest selective sweeps in the genome are all found around genes known to be important for resistance [6, 19]. Furthermore, a whole-genome scan of copy number variants (CNVs) in the Ag1000g data revealed that increases in gene copy number were highly enriched in clusters of detoxification genes, pointing to a potentially widespread mechanism for increased gene expression [20], which, in some cases, may elevate resistance. A number of gene duplications were observed around the *Cyp6aa/Cyp6p* gene family cluster on chromosome 2R, and the majority of these duplications included the gene *Cyp6aa1* (Figure 1). *Cyp6aa1* has been found to be overexpressed in pyrethroid resistant populations in congeneric species [21–23], but it has received very little attention compared to known insecticide-metabolizing genes such as *Cyp6m2*[24, 25], *Cyp6p3* [15] and *Cyp9k1* [26] and its importance in resistance in An. *gambiae* remains unknown.

**Figure 1.**
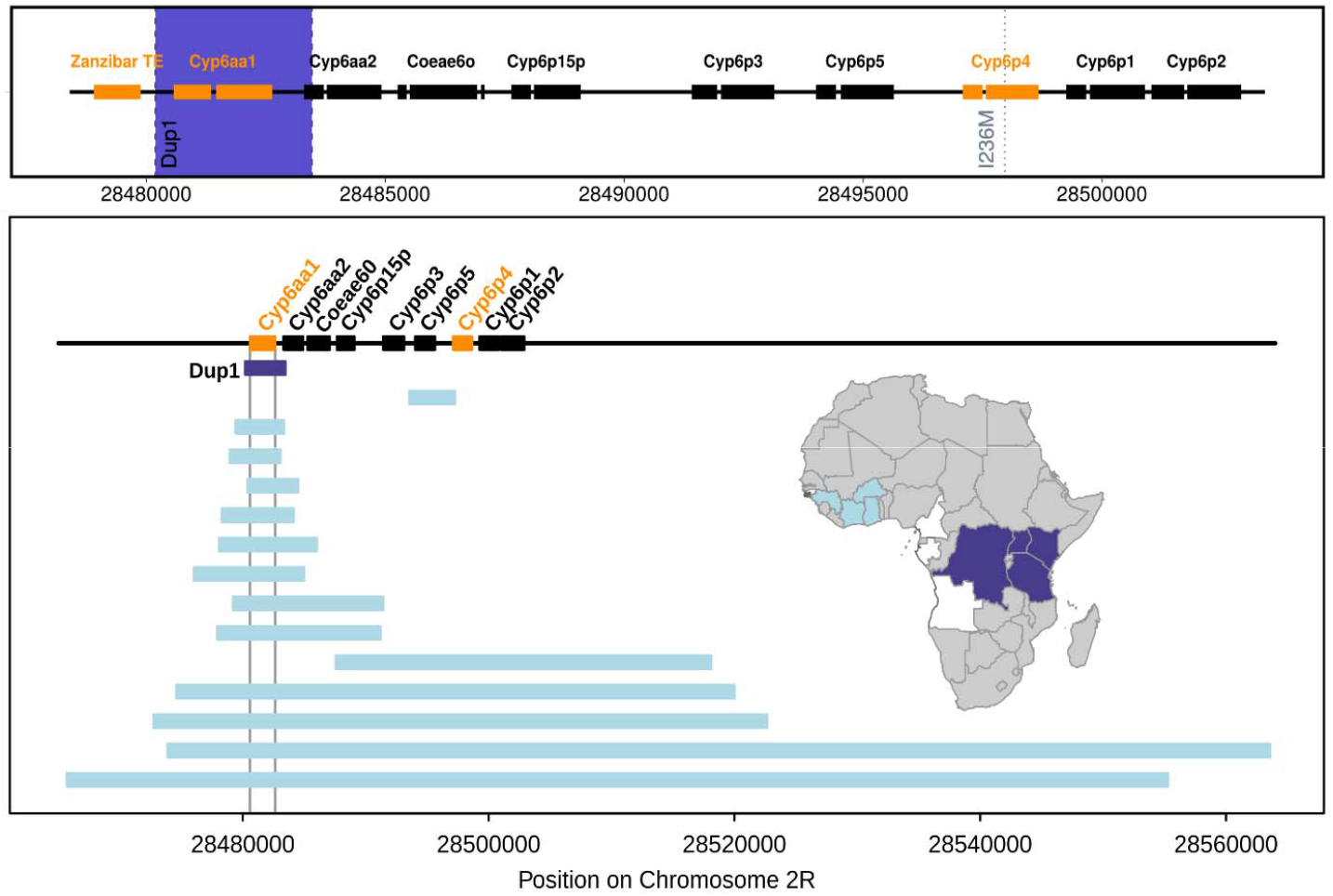
Schematic of the mutational events around the *Cyp6aa/Cyp6p* cluster on chromosome 2R in *Anopheles gambiae*. Upper panel shows the three mutational events observed in East and Central African samples. Orange indicates the genes/ transposable elements involved and the blue band the extent of the *Cyp6aap-Dup1* CNV. The lower panel, redrawn from [20], shows the genomic extent and geographic distribution of the CNVs that have been observed around the *Cyp6aa/Cyp6p* cluster, 13 out of 15 include *Cyp6aa1*. The dark blue shading on the map indicates the geographic extent of *Cyp6aap-Dup1*. Data from Uganda, Kenya and DRC are reported extensively in the main text, in addition 27.4% (n=84) *of Anopheles gambiae* females sampled from northern Tanzania in 2018 were triple mutant carriers.

In this study, we examine a strong selective sweep detected in the *Cyp6aa/Cyp6p* genomic region in samples of *An. gambiae s.s*. from Uganda and Western Kenya. We find that the sweep is closely associated with three mutations (a SNP in *Cyp6p4*, a duplication of *Cyp6aa1* and a partial transposable element insertion termed ZZB-TE) in tight physical and statistical linkage. The triple-mutant haplotype is associated with a high-level of pyrethroid resistance, most notably to deltamethrin. The three mutations appeared sequentially, leading to successive selective sweeps, with the triple-mutant haplotype replacing earlier variants and then spreading rapidly across East and Central Africa. We show that this haplotype is under positive selection and causes increased expression of key cytochrome P450s and through recombinant protein expression using both an *E. coli* and an *Sf9*-baculovirus system we show that both CYP6AA1 and CYP6P4 are capable of metabolizing pyrethroid insecticides.

## Methods

### Interrogation of the Ag1000g dataset and identification of tagging markers

Whole genome sequence data in the Ag1000g data set have previously revealed a strong selective sweep in Ugandan populations around the *Cyp6aa/Cyp6p* cluster [19] (Figure 1). An isoleucine to methionine substitution in codon 236 of CYP6P4 (see extended data Figure 10b in [19]) was identified provisionally as a swept haplotype tagging SNP. Previous work [20] had shown that a duplication of the *Cyp6aa1* gene was also observed in these samples (previously termed *Cyp6aap-Dup1*). To objectively determine how these mutations segregated with the observed selective sweep [19] we grouped the 206 Ugandan haplotypes (n=103 diploid individuals) by similarity using the 1000 SNPs located immediately upstream and downstream of the start of the *Cyp6aa/Cyp6p gene* cluster (500 non-singleton SNPs in each direction from position 2R:28,480,576). Distances were calculated with the *pairwise_distance* function in *scikit-allel* [27] and converted to a nucleotide divergence matrix (*Dxy* statistic) by correcting the distance by the number of sequencing-accessible bases in that region. We defined clusters of highly similar haplotypes by hierarchical clustering with a cutoff distance of 0.001. This resulted in the identification of a cluster of 122 highly similar haplotypes. To determine whether the haplotype cluster showed signs of a selective sweep, we estimated the extended haplotype homozygosity (EHH) decay within each of the haplotype groupings, around two different focal loci: (i) the putative sweep SNP marker *Cyp6p4-236M* (2R:28,497,967 +/- 200 kbp; total 14,243 phased variants), and (ii) the 5⍰ and 3⍰ breakpoints of the *Cyp6aap-Dup1* duplication (2R:28,480,189 - 200 kbp and 2R:28,483,475 + 200 kbp; total 14,398 phased variants). We used the *ehh_decay* function of *scikit-allel* [27].

We also used the haplotype groupings identified above to calculate the Garud *H* statistics [28] and the haplotypic diversity at the *Cyp6aa/Cyp6p* cluster locus (coordinates 2R:28,480,576 to 2R:28,505,816). Specifically, we used the *moving_garud_h* and *moving_haplotype_diversity* functions in *scikit-allel* to obtain a series of estimates for each statistic in blocks of 100 variants located within the cluster, and used a block-jackknife procedure to calculate averages and standard errors of each estimate (*jackknife* function in *scikit-allel misc* module). Python scripts to reproduce these analyses are available https://github.com/xgrau/cyp6-AgUganda together with genomic variation data https://www.malariagen.net/data/ag1000g-phase1-ar3.1.

### Molecular screening of colony and wild caught mosquitoes

Locked-nucleic acid (LNA) probe-based PCR diagnostics were designed for all three mutations (Supplementary materials Appendix 1).

Genotype:phenotype association testing was performed using two colonies of *An. gambiae s.s*., BusiaUG (resistant) and Mbita (susceptible). The BusiaUG strain was established in the lab in November 2018 from Busia, eastern Uganda and exhibits high resistance to pyrethroids and organochlorines (<10% mortality following WHO exposure test), but full susceptibility to organophosphates and carbamates (Oruni *et al*. unpublished). The Mbita strain was first colonised from Mbita Point, Kenya in 1999, and is fully susceptible to pyrethroids. Colonies were reared in insectaries targeted to 25-27°C and 70-80% relative humidity.

Freshly emerged females from the Mbita line were mated with 3-5 day-old BusiaUG males and then blood fed. The offspring from this cross were then crossed back to the parental BusiaUG line. This design was chosen as resistance variants are often recessive. The resultant backcrossed 3-5-day-old females were exposed for one hour to deltamethrin, permethrin or α-cypermethrin (the three insecticides most commonly used on LLINs) or DDT (a non-pyrethroid sodium channel antagonist), following WHO standard procedures. Mosquitoes were maintained on a 10% sugar solution after exposure and mortality was recorded 24 hours post exposure.

<u>Democratic Republic of Congo</u> *Anopheles gambiae s.s*. were obtained from the President’s Malaria Initiative supported entomological surveillance project [29] and from collections conducted by Lynd *et al* [30]. Mosquitoes were collected by human landing catch or pyrethrum spray collection from 15 locations between 2013 and 2018. Resistance-phenotyped individuals were also obtained from Pwamba, Bassa and Fiwa in Nord Ubangi Province in 2016. These mosquitoes were assessed for susceptibility to deltamethrin or permethrin using a standard WHO tube assay or a cone bioassay where Permanet 3.0 (deltamethrin plus PBO) or Olyset Plus (permethrin plus PBO) were the test nets [23]. <u>Kenya and Uganda.</u> Collection details for specimens from contiguous areas of western Kenya and eastern Uganda have been published previously [20, 24, 25]. <u>Tanzania.</u> Mosquito collections were conducted in Geita, Bagamoyo and Muleba districts of Tanzania in 2018.

Mosquitoes from all collections were genotyped at the ZZB-TE, *Cyp6p4-236M* and *Cyp6aap-Dup1* loci and genotype:phenotype association testing was performed by Fisher’s exact tests.

### Analysis of temporal change

The most complete time series of ZZB-TE, *Cyp6p4-236M* and *Cyp6aap-Dup1* allele frequencies were available from Kabondo, DRC; western Kenya and eastern Uganda. To model allele frequency changes over time we estimated the three parameters of the standard recursive population genetic model (allele frequency at time zero, selection coefficient and dominance coefficient) using a maximum likelihood approach assuming a binomial distribution: an approach previously applied to insecticide target-site resistance mutations [31]. The analysis was performed in R (http://www.r-project.org). An estimated generation time of one calendar month was used as in previous studies [31].

### Recombinant protein and insecticide metabolism

The CYP6P4-236M variant, and its redox partner cytochrome P450 reductase gene (CPR), were expressed in *E. coli* as per standard protocols [32] (Supplementary materials Appendix 2). Initial efforts to generate recombinant *CYP6AA1* in an *E. coli* system with optimised codon usage failed, and we therefore used an Sf9-baculovirus-based expression system. Since P450 catalytic activity is dependent on electrons supplied by NADPH via CPR, insecticide metabolism was assayed with cell pellets of CYP6P4/CPR or CYP6AA1/CPR in the presence or absence of NADPH. The depletion of the substrate and the appearance of metabolites were monitored by reverse-phase HPLC (Supplementary materials Appendix 3).

### Estimation of gene expression of key P450s in triple mutant haplotype

To determine whether the presence of the triple mutant haplotype type was associated with differential expression of genes, we examined individual females from the BusiaUg colony (Triple mutant Freq. 0.297; 95%CI 0.233-0.370). Two legs were removed from individual mosquitoes for DNA analysis with the remaining mosquito kept for RNA analysis. DNA was extracted from legs by boiling in STE buffer at 95°C for 90 minutes and individuals were genotyped using the LNA qPCR assays. RNA was then extracted individually from 8 mosquitoes in each genotypic group - homozygotes for the triple mutant haplotype, wild-type homozygotes, and heterozygotes, using the Arcturus Picopure RNA isolation kit (Thermofisher). We then performed SYBR green based qPCR to measure the expression of *Cyp6aa1* and *Cyp6p4* together with the known resistance-linked variant *Cyp6p3* using the housekeeping genes 40s ribosomal protein S7 (AGAP010592) and elongation factor Tu (AGAP005128) for normalisation. The ΔΔCT values were tested for normality and homogeneity of variances using the Shapiro-Wilks test, and the Bartlett test, respectively. A significant difference in gene expression between the genotypic groups was determined by a two-tailed two-sample Student’s t-test on ΔΔCT values, with a threshold of P=0.05.

## Results

Hierarchical clustering of 206 Ag1000g haplotypes from *An. gambiae s.s*. from Uganda resulted in identification of a cluster of 122 haplotypes around the *Cyp6aa/Cyp6p* gene cluster, putatively representing a swept haplotype in this region. To characterise the signatures of selection in Ugandan haplotypes around this cluster, we examined the profile of extended haplotype homozygosity around the position of the *Cyp6p4-236M* SNP and around the CNV in *Cyp6aa1*. In both cases, we found that the putative swept haplotype had longer stretches of homozygosity than wild-type haplotypes (Figure 2). In addition, we found that *An. gambiae s.s*. from Uganda had reduced haplotypic diversity along the entire *Cyp6aa/Cyp6p* gene cluster (*h* = 0.339746 +/- 0.005664 standard error) and a combination of Garud’s *H* statistics that was indicative of a hard selective sweep in this region (high *H_12_* = 0.821867 +/- 0.006308 SE; low *H_2_/H_1_* = 0.016779 +/- 0.000228 SE)[28]. These results confirm that the haplotypes we have identified have undergone a selective sweep. We then used iterative read mapping of individuals homozygous for the sweep to search for additional mutations that might be distinctive of the haplotype. This revealed that a partial copy of a Ty3/Gypsy Zanzibar transposon insertion (termed ZZB-TE), lacking functional open reading frames, was linked to *Cyp6p4-236M* and *Cyp6aap-Dup1* (Figure 2).

**Figure 2.**
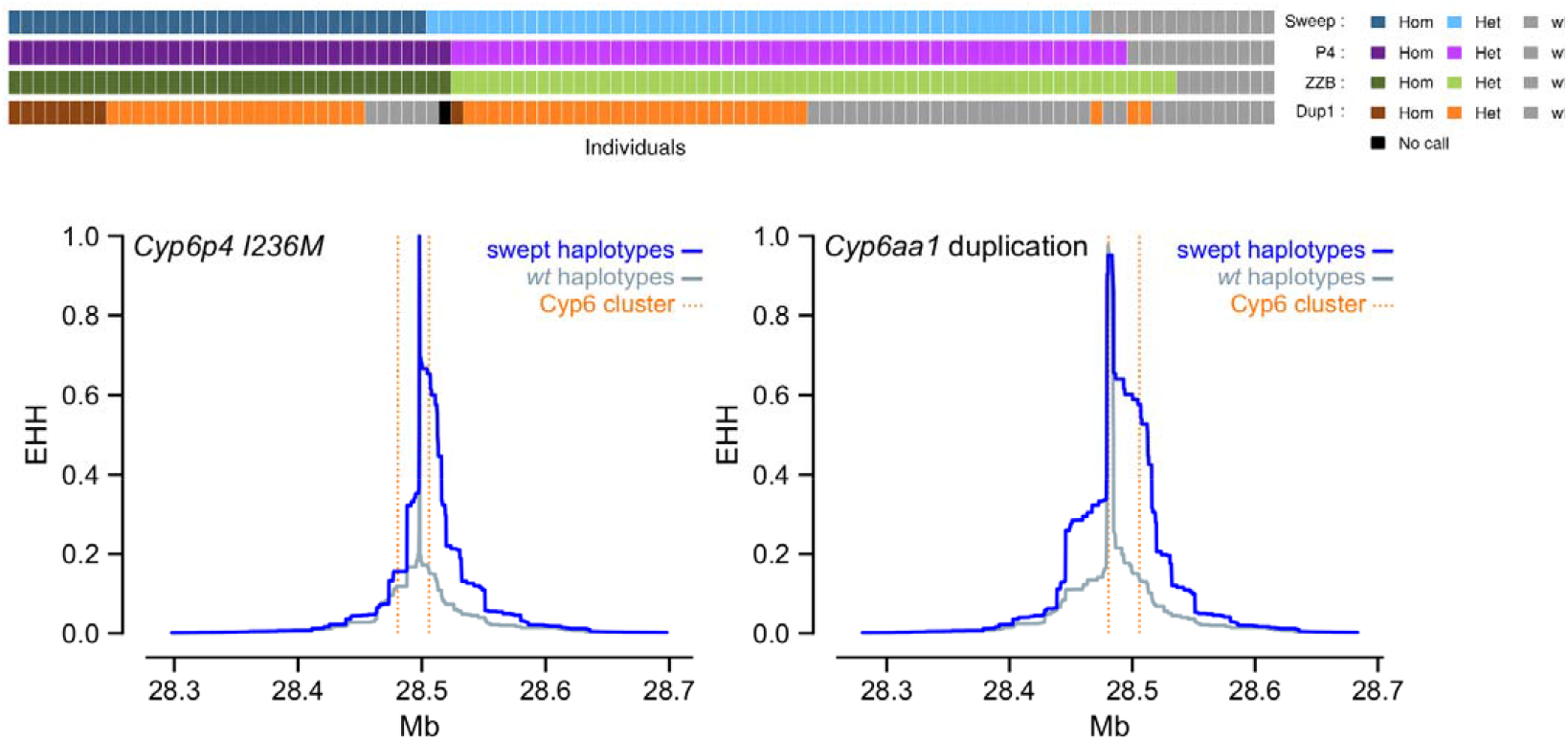
Selective sweep around the *Cyp6aa/Cyp6p* cluster in *Anopheles gambiae* from Uganda. Upper panel Haplotype contingency for the selective sweep (“Sweep”), *Cyp6p4-236M* SNP (“P4”) and the *Cyp6aap-Dup1* (“Dup1”). Each vertical bar represents a single haplotype, colour-coded to show whether it is a copy of the swept haplotype (blue) and whether it carries the SNP (purple), the TE insertion (green) and the duplication (orange). The swept haplotype and the *Cyp6p4-236M* SNP overlap almost completely, while the duplication is found on a subset of these haplotypes. Lower panel Extended Haplotype Homozygosity (EHH) plots around the *CYP6P4-236M* and *CYP6AA1-Dup1* variants show slower loss of homozygosity in the swept haplotypes than in the wild-type.

### The evolution of the ZZB-TE, *Cyp6p4-236M* and *Cyp6aap-Dup1* haplotypes

Based upon Ag1000g data and a time series of collections from Central and East Africa, we were able to trace the sequence of mutational events (ZZB-TE, *Cyp6p4-236M* and *Cyp6aap-Dup1*) and reconstruct the evolutionary history of the swept haplotype. Among the Ag1000g data [19], the *Cyp6p4-236M* mutation was only observed in collections from eastern Uganda (collected in 2012) suggesting that this mutation originated in the eastern Ugandan/western Kenyan region. In a screen of collections from Uganda and Kenya predating the Ag1000g collections by eight years (2004) (Fig 3) only the ZZB-TE insertion was detected, although the sample size was too small (n=4) to conclude that the *Cyp6p4-236M* allele was absent. The *Cyp6p4-236M* mutation was first observed in this region in 2005 (frequency *Cyp6p4-236M* =0.10) in individuals carrying the ZZB-TE mutation, whilst the *Cyp6aap-Dup1* CNV was first recorded in 2008 (proportion of individuals with Dup1=0.8%). This inferred sequence of events may explain why ZZB-TE and *Cyp6p4-236M* mutations are in tighter statistical linkage with each other than with *Cyp6aap-Dup1* (Figure 2), despite the closer proximity of ZZB-TE and *Cyp6aap-Dup1* (Figure 1). Given the very tight association between the ZZB-TE insertion and the *Cyp6p4-236M* SNP, we will henceforth refer to the *Cyp6p4-236M* (double mutant) haplotype and the *Cyp6aap-Dup1* (triple mutant). The double mutant haplotype shows a steady increase in frequency between 2004 and 2011 in Kenya (Figure 3); possibly in response to the introduction and subsequent intensification of bednet distribution programmes [29, 33, 34]. Following its appearance in 2008, the triple mutant haplotype, rapidly increased towards fixation in both collections from Uganda and Kenya, replacing the double mutant. This haplotype replacement and the observation that the triple mutant is the only non-wildtype haplotype observed outside Kenya/Uganda (such as in Tanzania and DRC, Figure 1 and 3) strongly implies an additional selective advantage to the triple mutant. The time series data from across DRC are particularly striking both in terms of the speed of increase of the triple mutant but also the north-south heterogeneity, with very low frequencies in the more southerly provinces (Figure 3).

**Figure 3.**
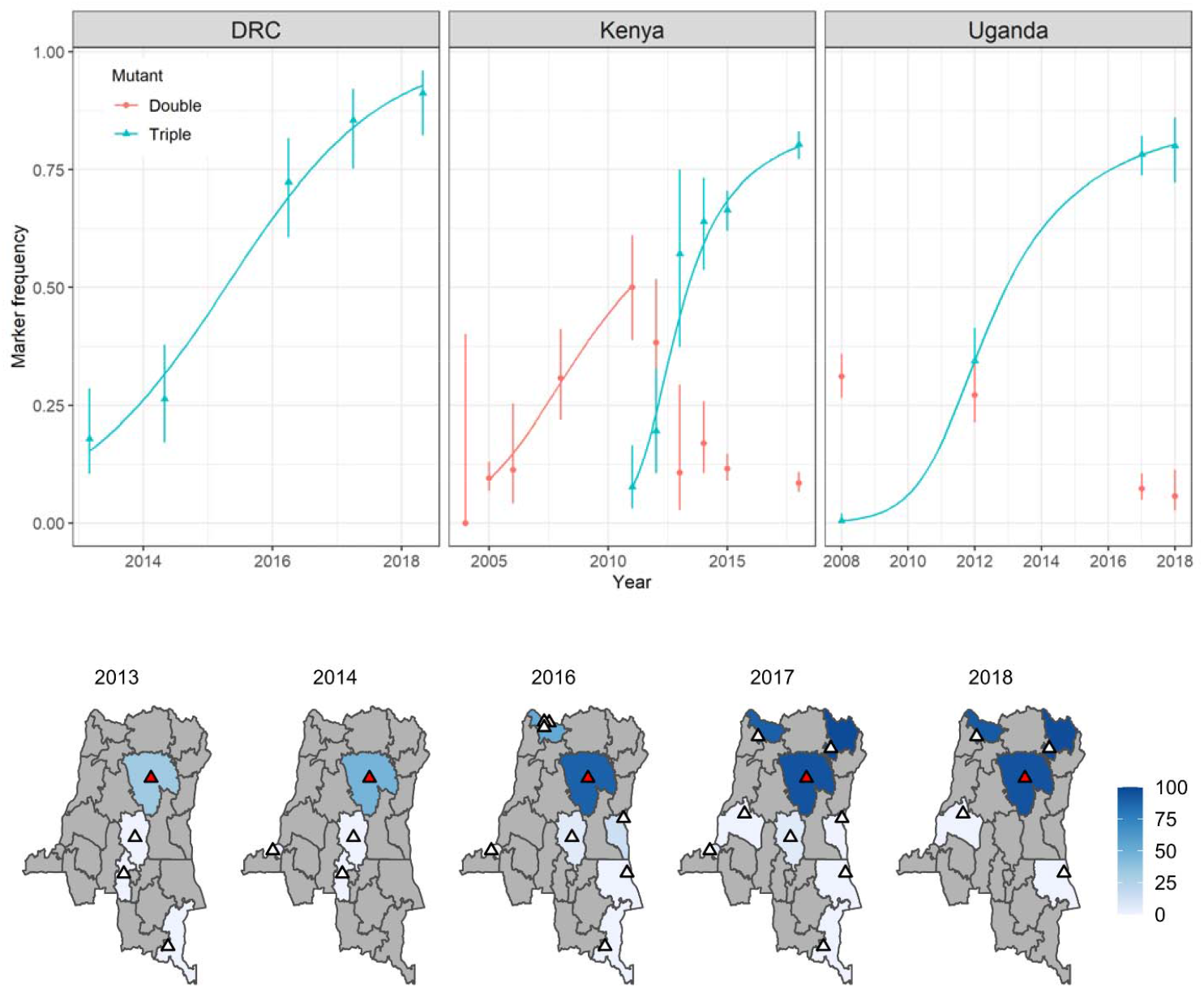
Observed and modelled changes in *Cyp6aa/Cyp6p* haplotype frequencies over time in *Anopheles gambiae s.s*. populations from DRC, Kenya and Uganda. Upper panel shows mutation frequency estimates derived from wild caught individuals from Kabondo, DRC; western Kenya and eastern Uganda, The 95% CIs for each observed data point were calculated according to[47]. Expected data generated from simultaneous maximum likelihood estimates of initial frequency and selection and dominance coefficients which were then used to parameterize standard recursive allele frequency change equation. The lower panel shows data from DRC tracking the emergence and spread of the triple mutant haplotype. Triangles indicate collection locations within each district. The red triangle shows the Kabondo sample site for which there was the most complete time series.

### Identifying potential drivers of haplotype frequency increase

The genomic region into which the ZZB-TE inserted does not show histone signals of regulatory activation (H3K27ac, H3K9ac) or repression (H3K9me3), and ATAC-seq data suggests it is not in an open chromatin region [35]. However we took two in silico approaches to determine whether the ZZB-TE insertion (748bp) carried putative regulatory variants. The inserted region had 98.5% sequence identity to a putative enhancer (2R:45966598-45966822) identified by homology with *Drosophila melanogaster* [35]. However, despite the similarity, the insertion lacked enhancer-like combination of chromatin marks identified in [35] and its potential regulatory role in nearby genes is unclear. The second approach involved screening the ZZB-TE inserted sequence for putative enhancers using iEnhancer-2L[36] and iEnhancer-EL [37]. In a windowed analysis of 200bp with a 1bp step across the entire length of ZZB-TE both predicted that some of the windows would have strong enhancer activity, however the windows were not-concordant, precluding further analysis.

Given that cytochrome P450 mediated resistance is commonly associated with differential gene expression we performed transcription studies within the *Cyp6aa/Cyp6p* cluster between the most contrasting haplotypes, wild-type and triple mutant, present in the BusiaUg colony. The group homozygous for the triple mutant haplotype significantly overexpressed both *Cyp6aa1* (2.23-fold, 95% CI: 1.73-2.90, P=0.0003) and *Cyp6p4* (2.57-fold, 95% CI 1.25-5.93, P=0.039) compared to wild-type individuals. The ratio of expression of *Cyp6aa1* broadly reflected the expected pattern based on genotype (ie 2:1.5:1 for triple mutant homozygotes: heterozygotes: wild type genotypes, respectively. Figure 4). As a control we examined a neighbouring, very commonly resistance-associated gene, *Cyp6p3*, but triple mutant and wild type homozygotes did not differ significantly in expression (1.33 fold, 95% CI 0.64-2.74, P>0.05).

**Figure 4.**
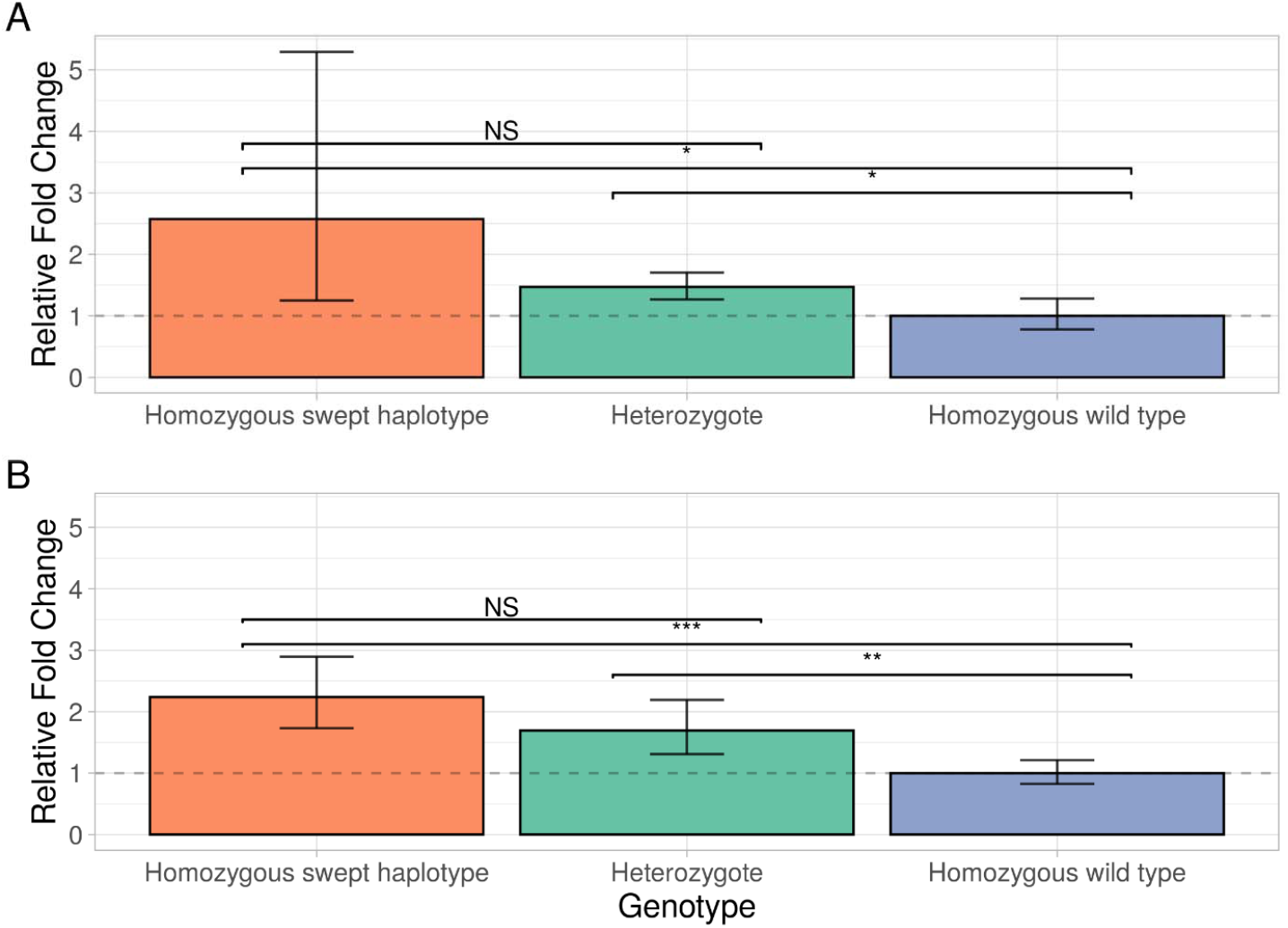
Gene expression analysis of *Cyp6p4* and *Cyp6aa1* in genotyped *Anopheles gambiae* females. Relative fold change for *Cyp6p4* (Panel A) and *Cyp6aa1* (Panel B), comparing individuals homozygous for the swept triple mutant haplotype and heterozygotes in the BusiaUG colony to wild-type individuals. 95% confidence intervals are shown. Asterisks indicate statistical significance in two-tailed students t-tests; *** p ≤ 0.001, ** P ≤ 0.01, ** P ≤ 0.05.

To investigate whether resistance may be driven at least in part by an effect of the allelic variant on metabolic activity of CYP6P4, we expressed the wild-type (236I) and mutant (236M) forms in an *E. coli* based recombinant protein system (Supplementary materials Appendices 2 and 4). Both alleles were shown to be capable of metabolizing class I (permethrin) and II (deltamethrin) pyrethroids but there was no evidence that the mutant (236M) or wildtype (236I) alleles had different rates of pyrethroid depletion. We also expressed the duplicated P450 CYP6AA1, in an *Sf9*-baculovirus protein expression system. Again, metabolism assays demonstrated that the enzyme was capable of metabolizing both deltamethrin and permethrin (Supplementary materials Appendices 3 and 4). Depletion of deltamethrin was 36.6% greater (SE= 3.79) in the presence of NADPH than in the control (t-test: t=-9.67; d.f. = 8; P= 9.6 x10^-6^), demonstrating that CYP6AA1/CPR is capable of metabolizing deltamethrin *in vitro*. Similarly, permethrin was metabolised by CYP6AA1/CPR, with permethrin being depleted by 22.4% (SE = 0.63) compared to the control without NADPH (t-test: t= −31.08; d.f. = 14; P= 2.55 x10^-14^).

Given clear evidence of increased expression of both *Cyp6aa1* and *Cyp6p4* in the triple mutant haplotype and the ability of both enzymes to metabolise pyrethroids *in vitro*, we investigated whether the mutations were significantly associated with resistance *in vivo*. Exposure of *An. gambiae* females from Busia, Uganda and Nord Ubangi, DRC to new LLINs in cone assays resulted in negligible mortality to the pyrethroid only LLINs, Olyset and Permanent 2.0 (Figure 5). Simultaneous exposure to pyrethroid plus the P450 inhibitor PBO in Olyset + and the top of Permanent 3.0 nets resulted in a marked reduction in resistance, demonstrating that the resistance phenotype is substantially mediated by P450s. We performed laboratory backcrosses between additional mosquitoes from Busia with the pyrethroid susceptible Mbita colony, and found that the triple mutant haplotype was significantly associated with resistance to the most commonly used type II pyrethroids in LLINs: deltamethrin (Fisher’s exact test p=3.2×10^-6^) and alphacypermethrin (Fisher’s exact test p=5.9×10^-7^) resistance although not to permethrin (Fisher’s exact test p=0.06) nor, as a control, DDT (Fisher’s exact test p=0.84) (Table 1) in WHO tube assays. Similarly, specimens collected in 2016 from the DRC showed a strong association between the triple mutant genotype and survival rate 24 hours post-exposure to either 0.05% deltamethrin for 1 hour or 3-minute exposures to deltamethrin-treated sides of a new PermaNet 3.0 net (Table 2). No association was found in samples exposed to permethrin (24 hour WHO tube assay) or permethrin-treated Olyset Plus nets (3-minute WHO cone assay) (Table 2). Complete linkage of the three mutants in the BusiaUG colony and the DRC wild caught collections precludes determination of the relative contribution of each of the three mutations to the resistance phenotype but taken together these results demonstrate a strong impact of the triple mutant on the efficacy of pyrethroid resistance.

**Figure 5.**
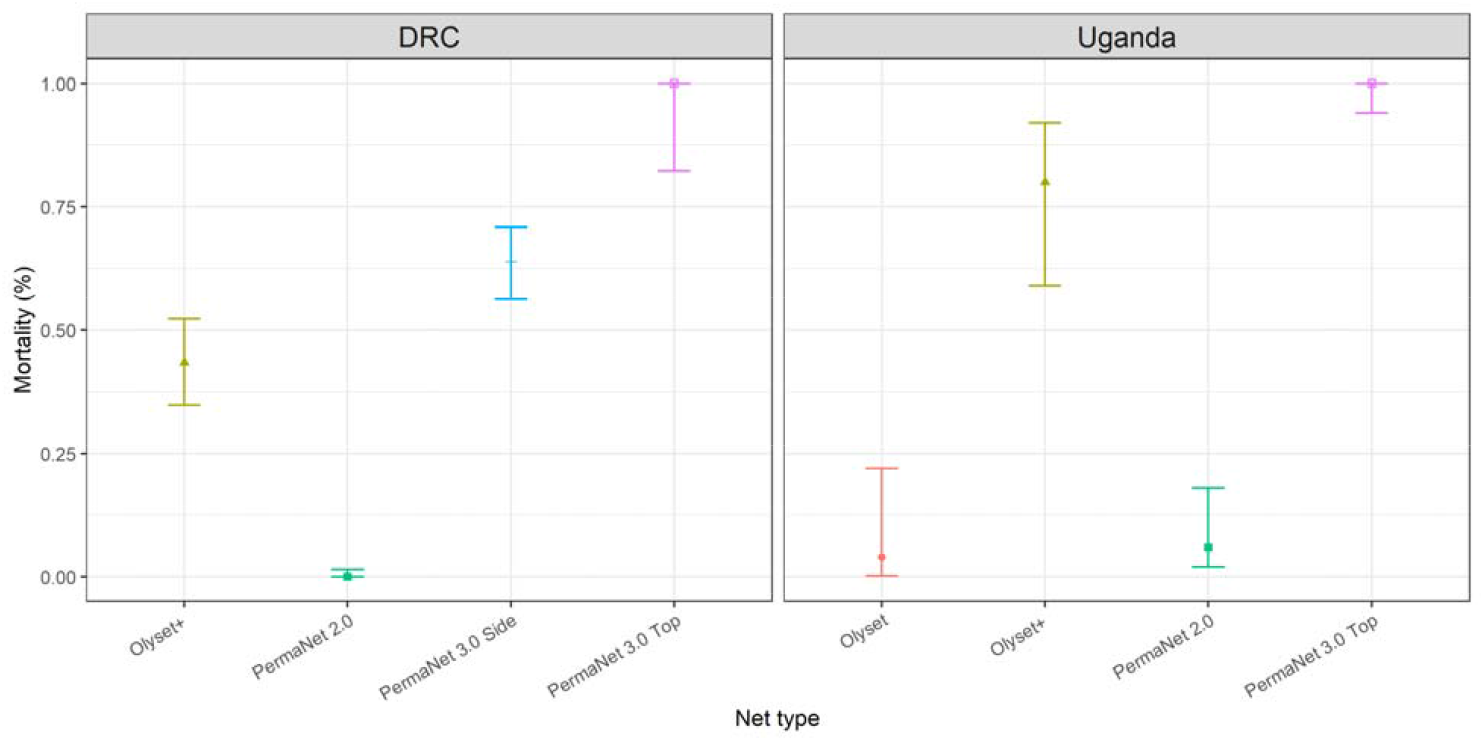
Patterns of insecticide resistance in *Anopheles gambiae* following exposure to permethrin and deltamethrin treated nets. Female *Anopheles gambiae s.s*. from the Democratic Republic of Congo (DRC) and Uganda exhibited very high resistance (low % mortality) in WHO cone assays to permethrin (Olyset) and deltamethrin (PermaNet 2.0) LLINs. There was an increase in mortality following exposure to an increased concentration of deltamethrin (PermaNet 3.0 Side) and very high mortality following exposure to pyrethroid plus the P450 inhibitor piperonyl butoxide (PBO) Olyset+ and PermaNet 3.0 Top. 95% confidence intervals are shown.

**Table 1.** Association between insecticide susceptibility as determined by WHO tube bioassay and triple mutant genotype in female *Anopheles gambiae* from a cross between BusiaUG and Mbita colonies. ^a^Odds ratios for allelic test and associated 95%CI

| **Insecticide** | **Phenotype** | ***Wildtype***<br>***homozygote*** | ***Heterozygote*** | ***Triple mutant***<br>***homozygote*** | **Test and p value** |
| --- | --- | --- | --- | --- | --- |
| **α-Cypermethrin**<br>**n=146** | **Alive** | 1 | 32 | 24 | Genotypic p=5.92x10<sup>-7</sup> |
|  | **Dead** | 14 | 68 | 7 | Allelic p=6.34x10<sup>-5</sup>; OR<sup>a</sup><br>2.74 (1.63-4.69) |
| **Deltamethrin**<br>**n=60** | **Alive** | 0 | 0 | 8 | Genotypic p=3.21x10<sup>-6</sup> |
|  | **Dead** | 15 | 26 | 11 | Allelic- Cannot be<br>calculated |
| **Permethrin**<br>**n=53** | **Alive** | 1 | 18 | 7 | Genotypic p=0.059 |
|  | **Dead** | 5 | 20 | 2 | Allelic p=0.08; OR 1.99<br>(0.86-4.67) |
| **DDT**<br>**n=65** | **Alive** | 0 | 14 | 1 | Genotypic p=1 |
|  | **Dead** | 3 | 45 | 2 | Allelic p=0.84; OR 1.19<br>(0.48-2.94) |

**Table 2.** Association between insecticide susceptibility as determined by WHO tube bioassay or WHO net bioassay and triple mutant genotype in wild-caught, female *Anopheles gambiae* from three locations in Nord Ubangi Province, Democratic Republic of Congo. ^a^Odds ratios for allelic test and associated 95%CI

| Site | Insecticide | Bioassay | Phenotype | *Wildtype*<br>*homozygote* | *Heterozygote* | *Triple*<br>*mutant*<br>*homozygote* | Test and p value |
| --- | --- | --- | --- | --- | --- | --- | --- |
| Fiwa | Permethrin<br>n=161 | Tube | Alive | 5 | 42 | 30 | Genotypic p=0.63 |
|  |  |  | Dead | 8 | 49 | 27 | Allelic p=0.42; OR<sup>a</sup> 1.23 (0.76-2.00) |
| Fiwa | Deltamethrin<br>n=78 | Tube | Alive | 6 | 14 | 18 | Genotypic p=6.21x10<sup>-3</sup> |
|  |  |  | Dead | 7 | 27 | 6 | Allelic p=0.04; OR 2.01 (1.01-4.06) |
| Bassa | Permethrin<br>n=95 | Tube | Alive | 1 | 19 | 18 | Genotypic p=0.79 |
|  |  |  | Dead | 2 | 32 | 23 | Allelic p=0.63; OR 1.21 (0.61-2.43) |
| Bassa | Deltamethrin<br>n=47 | Tube | Alive | 0 | 3 | 16 | Genotypic p=4.02x10<sup>-3</sup> |
|  |  |  | Dead | 1 | 16 | 11 | Allelic p=5.71x10<sup>-3</sup>; OR 5.44 (1.41-31.28) |
| Pwamba | Deltamethrin<br>n=40 | Tube | Alive | 7 | 9 | 7 | Genotypic p=0.066 |
|  |  |  | Dead | 9 | 9 | 1 | Allelic p=0.02; OR 2.93 (1.07-8.39) |
| Fiwa | Olyset Plus<br>n=106 | Net | Alive | 6 | 21 | 30 | Genotypic p=0.82 |
|  |  |  | Dead | 6 | 15 | 28 | Allelic p=0.88; OR 0.93 (0.49-1.77) |
| Fiwa | Permanet 3.0<br>n=155 | Net | Alive | 5 | 26 | 24 | Genotypic p=5.61x10<sup>-4</sup> |
|  |  |  | Dead | 30 | 51 | 19 | Allelic p=1.4x10<sup>-4</sup>; OR 2.56 (1.54-4.31) |

## Discussion

We have identified a sequential series of fitness-augmenting mutations in *An. gambiae* culminating in a triple-mutant haplotype with a large effect on pyrethroid resistance and which is spreading rapidly across East and Central Africa. The mutation that is probably the oldest in this series, the insertion of ZZB-TE, was first detected in 2004 in the malaria-endemic area around Lake Victoria, with the *Cyp6p4-236M* SNP evident in 2005 samples and the third, a duplication in *Cyp6aa1* detected in 2008. In samples collected only five years later, the triple mutant was detected hundreds of kilometres away in the DRC. These patterns suggest both a large fitness advantage arising from the triple mutant, and a frightening speed at which resistance-conferring mutations are able to spread within and across populations.

Second generation nets treated with the synergist PBO were shown to be much more effective than conventional nets in killing mosquitoes in populations where the triple mutant haplotype is present (Figure 5). Therefore the use of PBO bednets should be prioritised in the regions where this mutation is present. A strong corollary of this finding comes from the cluster randomised control trial conducted in Uganda, where the mutation is at high frequency, which demonstrated that malaria parasite prevalence in children <10 years old (12% vs 14%; Prevalence ratio = 0·84, 95% CI 0·72–0·98; P=0·029) and mean number of mosquitoes (Density ratio=0·25, 95% CI 0·18–0·3; P <0·0001) per house were significantly lower in villages that had received PBO LLINs relative to standard LLINs [4]. There is some evidence that the haplotype may be less strongly associated with resistance to permethrin than deltamethrin (and perhaps also alphacypermethrin), although both pyrethroids were metabolised by *Cyp6aa1* and *Cyp6p4*.

Our results highlight the importance of gene duplications for the evolution of insecticide resistance. In *An. gambiae*, duplications have recently been shown to be concentrated in regions associated with metabolic resistance, and over 40 such duplications have been described across the genome [20]. Thirteen different duplications have so far been described that encompass *Cyp6aa1* (Figure 1), both in West and East Africa and in the two sister-species *An. gambiae* ss and *An. coluzzii* [20]. It seems likely that these other *Cyp6aa1* duplications are also associated with pyrethroid resistance. For example, in *An. coluzzii* sampled from a highly insecticide-resistant population from Cote d’Ivoire, greater than 95% of individuals had Cyp6aap duplications (Supplementary materials Appendix 5). Five different duplications were observed in the collections from 2012 and 2017 and, whilst we detected no association with pyrethroid resistance, two duplications (*Cyp6aap-Dup7* and *Cyp6aap-Dup14*) showed significant increases in frequency over time. Moreover the total number of CNVs per sample (measured as presence of each of the 5 duplications, summed for each sample) increased significantly from an average of 1.59/ individual in 2012 to 2.08 in 2017 (Mann Whitney U test; P<0.0001); the mean number of duplications >2 indicates that there are multiple CNVs on the same haplotype. Duplications of the *Cyp6aa1* orthologue have been found in another important malaria vector, *An. funestus*, from west and central Africa [21]. The *Cyp6aa1* orthologue in *An. funestus*, which shares an 87% identity with *An. gambiae*, was also observed to metabolize permethrin and deltamethrin and, when expressed in transformed *Drosophila*, was associated with significant increases in resistance to both permethrin and deltamethrin relative to control mosquitoes [21]. This evolution of multiple *Cyp6aa1* duplications suggests this is an important Africa wide resistance mechanism

There are now several cases of insecticide resistance evolution where an initial mutation in a genomic region is followed by the spread of additional mutations on the resistant haplotype background [14, 38–42]. It is not yet clear whether the sequence of mutations that we have identified rely on each-other for effect, and thus could only have spread sequentially, or whether each additional mutation coincidentally appeared on the background of an already common mutant haplotype. In the case of *Cyp6p4-236M* and *Cyp6aap-Dup1* it seems that the latter is more likely, although we cannot exclude the possibility that the duplication affects the regulation of *Cyp6p4*. In contrast, the putative enhancer inserted with the ZZB transposon may affect either or both of *Cyp6p4* and *Cyp6aa1*, and may thus interact with the other two mutations in ways that have yet to be determined. Transposable elements can sometimes affect gene expression of neighbouring genes [38] and are abundant in mosquito genomes [43, 44]. Interestingly, in the common house mosquito, *Culex pipiens*, TEs have been found in the flanking regions of the *Ester* locus, a genomic region in which many independent gene duplications have arisen and spread worldwide in response to selection from organophosphates [45]. Clearly, both TEs and gene duplications are an understudied, yet common source of variation that may have important implications for vector control efforts. The appearance and rapid spread of the three mutations described here is broadly coincident with the scale up of LLIN coverage in DRC, Kenya and Uganda. The haplotype is a strongly predictive marker of high-level resistance to pyrethroids, is easily screened with a single diagnostic assay and we suggest should be used for both insecticide resistance monitoring strategies and for informing LLIN selection.[6, 29, 46].

## Acknowledgements

This work was supported by the National Institute of Allergy and Infectious Diseases ([NIAID] R01-AI116811) with additional support from the Medical Research Council (MR/P02520X/1). The latter grant is a UK-funded award and is part of the EDCTP2 programme supported by the European Union. HN and AO were supported by Wellcome Trust MSc Training Fellowships in Public Health and Tropical Medicine. SI is funded by the U.S. President’s Malaria Initiative. CM was supported by the Bill and Melinda Gates Foundation through the WHO (Award #54497) and by the Biovision Foundation (Grant No.BV HH-07). JLB is supported by DELTAS Africa Initiative grant # DEL-15-011 to THRiVE-2. The DELTAS Africa Initiative is an independent funding scheme of the African Academy of Sciences (AAS)’s Alliance for Accelerating Excellence in Science in Africa (AESA) and supported by the New Partnership for Africa’s Development Planning and Coordinating Agency (NEPAD Agency) with funding from the Welcome Trust grant # 107742/Z/15/Z and the UK government. SI is funded by the U.S. President’s Malaria Initiative. The findings and conclusions in this paper are those of the authors and do not necessarily represent the official position of CDC, the NIAID, the National Institutes of Health or the other donors.

## Author contributions

HN, AvH, MJIP, DW and MJD designed the study; HN, AvT, AO, DP, SCN, AL conducted lab and insectary experiments; HN, AvH, SCN, AL, ERL, ST, XG-B, DM, MJD performed analysis; AO, AL FW, EM, FA, SI, BK, DW, CM conducted field collections and phenotyping; HN, ERL, DW, MJD wrote the manuscript with input from all authors; JLB, CM, MJIP, DW and MJD supervised the study; all authors approved the final version of the manuscript.

## Notes

### Competing Interest Statement

The authors have declared no competing interest.

### Summary of Updates

Some small changes following internal review

