## Supplementary material for "Identification of a rapidly-spreading triple mutant for high-level metabolic insecticide resistance in *Anopheles gambiae* provides a real-time molecular diagnostic for anti-malarial intervention deployment": Supplmentary data

### Supplementary materials

#### Appendix 1. LNA assay for haplotype variant detection.

Three, independent, locked-nucleic acid (LNA) probe-based PCR assays were designed to genotype the *Cyp6p4-I236M* SNP*,* the insertion of the partial ZZB-TE, and the duplication of *Cyp6aa1.* For the *Cyp6p4*-I236M assay, LNA-probes labelled with Fam and Hex target the wild type and mutant alleles respectively, whilst ZZB-TE locus is targeted by Hex and Fam probes complementary to the wild type DNA and the insert (mutant) DNA respectively. *Cyp6p4*-I236M and ZZB-TE assays produce codominant markers. The duplicated region of the haplotype, *Cyp6aap*-Dup1, is detected using a combination of three primers and three probes. Primers produce two amplicons, one within the duplicated region and one spanning the junction site of the duplication. A Hex-labelled probe targets the duplicated region, a Cy5-labelled probe is complementary to DNA external to the duplication to provide a reference value for single copy gDNA and a Fam-labelled probe hybridizes to the junction between duplicated genes to detect mutant duplication genotypes. Wild-type homozygotes are indicated by the absence of Fam signal in the presence of both Cy5 and Hex. Heterozygotes and homozygotes for the duplication are differentiated by analysis of the ratio of the Hex, Fam and Cy5 Ct values: 2x Cy5 - (Fam + Hex). Ratio values are then arranged in ascending order and plotted as a line graph. Heterozygotes and homozygotes can be differentiated by a change in the gradient of the line

All LNA probes were designed (http://biophysics.idtdna.com/) with a Tm difference of at least 16°C between target and non-target sequence. (Supplementary materials Table 1). PCR reactions were carried out in a total reaction volume of 10µl, consisting of 1X Luna® Universal Probe qPCR Master Mix (New England Biolabs, USA), 0.35µM of each primer (Supplementary materials Table 1), 0.2µM of each probe and 1µl of genomic *Anopheles* DNA. Reaction conditions consisted of a 3 minute denature at 95°C; 40 cycles of denaturation for 10 seconds at 95°C, annealing for 20 seconds at 58°C for the Zanzibar-6P assay, or 57°C for the *Cyp6aa1*-Dup1 assay, followed by an extension of 10 seconds at 72°C. The *Cyp6p4*-I236M assay consisted of a 3 minute denature at 95°C; 20 cycles of denaturation for 15 seconds at 95°C, annealing for 30 seconds at 66°C; 23 cycles of denaturation for 10 seconds at 95°C, annealing for 20 seconds at 58°C, and an extension of 10 seconds at 72°C. Reactions were run on an AriaMx Real-Time PCR system (Agilent, USA) equipped with Fam, Hex and Cy5 filters.

| Assay | Primer/Probe | 5'modification | Sequence | 3'modification |
|---|---|---|---|---|
| *Cyp6p4*-I236M | CYP6P4_i236M_F | | AGTTTATGTTTGCGACCACGTT | |
| | CYP6P4_i236M_R | | TCCACCGTCTCGCGCACAAC | |
| | CYP6P4_I-wild | FAM | TTC+ATGC+C+G+ATGC | Iowa Black®FQ |
| | CYP6P4_M-Mutant | HEX | TTC+ATGC+C+C+ATGC | Iowa Black®FQ |
| ZZB-TE | ZZB_flank_A | | CAAAATCAATGKCACRGAGC | |
| | ZZB_flank_B | | CGCTACAATGAAKGGAAAGTC | |
| | ZZB_int_A | | CATTACATGGCGACCGTACCT | |
| | ZZB_MUT | FAM | A+C+ATTA+CA+CTTTGT+CA+GTA | Iowa Black®FQ |
| | ZZB_WT | HEX | GATG+TT+CTTTK+T+CA+G+TATT | Iowa Black®FQ |
| *Cyp6aap*-Dup1 | AA1_Dup1_ins1 | | CAGTGCGGTACGCTCGTTAA | |
| | AA1_Dup1_outs | | GGATCGGTTTACAGCGGACG | |
| | AA1_Dup1_ins2 | | CATCACCTGTGCTCGCAARTT | |
| | AA1_Dup1_outs | Cy5 | CCAT+CA+C+CGAA+CG | Iowa Black®Rq-Sp |
| | AA1_Dup1_ins | HEX | AGAA+CCTGCA+C+CAA | Iowa Black®FQ |
| | AA1_Dup1_junct | FAM | A+CA+AT+TAATT+G+CAT+CGG | Iowa Black®FQ |

**Supplementary table 1.1. Primer and probe sequences for the LNA assays to detect the *Cyp6* cluster variants.**

### Appendix 2. Heterologous expression of *An. gambiae* CYP6P4 and CPR using an *E.coli* expression system

#### Synthesis and site directed mutagenesis

Sequences of CYP6P4 variant 1 (236M) were synthesised by Eurofins Genomics. For optimal expression of the variants in bacteria, the gene sequences were modified to contain *E. coli* optimized codons and a bacterial *ompA* leader sequence upstream of the start codon. The leader sequence had an Nde1 restriction site at the start codon while its terminal end was linked to the 5' end of the cytochrome P450 gene with an alanine-proline linker sequence. *Cyp6p4* variants were cloned into a pCY066 plasmid.

The reference sequence (PEST strain) *Cyp6p4*-236I was generated by site directed mutagenesis (SDM). The mutagenic primers were designed in NEBaseChanger and a NEB Q5® site directed mutagenesis kit used for SDM. We designed the mutagenic primers such that the 5' ends of the forward and reverse primers would anneal back to back on the target site to amplify the gene as well as the plasmid. Apart from the single nucleotide change, the primers also introduced restriction sites used for screening SDM mutants. SDM, screening and sequencing primers are shown on Table 2.1. SDM was carried out as per the protocol provided by the manufacturer. The protocol had three steps. The exponential amplification step to introduce mutations, the Kinase, ligation and Dpnl treatment step to recyclize the plasmid and the transformation step to introduce plasmids in bacteria for amplification and screening. Agar plates, supplemented with ampicillin (50mg/ml) were used to select transformed bacteria. Screening successful SDM mutants was approached in two ways, a nested PCR that comprised of colony PCR to confirm the colonies had the insert and a second PCR targeting the mutagenized region. Sequencing was performed to confirm successful SDM. The other approach was restriction analysis where single colonies from the selection plate were cultured overnight in Luria-Bertani broth at 37°C and their plasmids purified using a QIAprep® Spin Miniprep kit as per manufacturer's protocol. Restriction analysis with Nde1 and Not1 was done to screen colonies with an intact gene and plasmid sequences. Another digest with enzymes targeting the restriction sites introduced alongside the single nucleotide change was done. In all restriction analysis, 5 µl of plasmids and 5 µl of restriction master mix were incubated at 37°C overnight and visualised by gel electrophoresis. The restriction master mix contained 0. 2 µl of restriction enzyme, 1 µl of NEB buffer 3 for Nde1 and Not1 or NEB cutsmart buffer for Bmt1, Stu1 and BsrG1, and topped up with molecular water to 5 µl. Sequencing was done to confirm the screened successful SDM mutants.

| Activity | Orientation | Sequence |
|---|---|---|
| **SDM primers** | | |
| 6P4_V1 to 6P4_REF | Forward | aacgcatcGGCATGAAATTAACGGATG; |
| | Reverse | tggctagcCCTTTAAAGGTCGTAGCAAAC |
| **Screening and sequencing primers** | | |
| 6P4 Mutants | Forward 1 | ACGCGAAAGCCGATCCACTGA |
| | Forward 2 | CGGGAAATGAACAACGTACAA |
| | Forward 3 | ACCGTTCCGGGTACCAAACAT |
| | Forward 4 | CATTGCGATTGCGGTAGCCTTA |
| | Forward 5 | GCTATGTCCTTACCGCATTCGT |
| | Reverse 1 | GAAATTCCCCAGCGACCTGTTG |
| | Reverse 4 | GGTTGACGGGGAAAGGATGAAG |
| | Reverse 5 | ACGTAACAGCGTAATCAGTCCG |

***Table 2.1. Back to back mutagenic and screening primers used for site directed mutagenesis***

#### Transformation

Each variant was co-transformed with *An. gambiae* P450 reductase (pACYC-AgCPR)[1] in Stellar™ competent *E. coli* cells (Clontech). Fifty microliters of cells thawed on ice were aliquoted into 15 ml loose-cap tubes where 1 µl of the variant and 1 µl of pACYC-AgCPR plasmids were added. The cells were incubated on ice for 30 minutes, heat shocked in a 42°C water bath for 20 seconds and further incubated on ice for 5 minutes. Into the cells, 500 µl of pre-warmed LB broth was added followed by incubation at 37 °C in a shaking incubator set at 200 revolutions per minute (rpm) for 1 hour. The cells were plated in LB agar plates supplemented with 50mg/ml ampicillin and 25 mg/ml chloramphenicol and further incubated overnight at 37°C to select cells that were transformed with both plasmids.

#### P450 expression

A few colonies from the selection plate were inoculated in 8 ml LB broth supplemented with 8 µl of 50mg/ml ampicillin and 8 µl 25 mg/ml chloramphenicol. The cells were cultured at 37°C in a shaking incubator set at 200 rpm for 12 hours to generate an 8 ml starter culture . Three mls of starter culture used to inoculate 200 ml modified terrific broth supplemented with 50 µl of 4000X trace elements, 200 µl of 1M thiamine hydrochloride, 200 µl of 50mg/ml ampicillin and 200 µl of 25 mg/ml chloramphenicol in a 1 litre Erlenmeyer flask. To prepare 100ml 4000x trace elements stock, 2.45 ferric citrate was added in about 50ml distilled water, stirred over moderate heat until dissolved and then left to cool. To the ferric citrate solution, 10 ml concentrated HCl, 0.131 g of $ZnCl_2$, 0.2 g of $CoCl_2{\cdot}6H_2O$, 0.2 g of $Na_2MoO_4{\cdot}2H_2O$, 0.1 g of $CaCl_2{\cdot}2H_2O$, 0.127 g of $CuCl_2{\cdot}2H_2O$, and 0.05 g of $H_3BO_3$ were added. The mixture was stirred until dissolved and topped up to 100ml with distilled water. The cells were cultured at 37°C in a shaking incubator set at 200 rpm and their optical density (OD) monitored periodically. At OD 0.4, 200 µl of 1M δ-Aminolevulinic acid (ALA), a heme precursor was added to the cultures. The incubation temperature and shaking speed were adjusted to 21°C and 120 rpm respectively. The culture monitored periodically until the OD reached 0.6 when expression of P450 was induced with 1mM Isopropyl-β-D-thiogalactopyranoside (IPTG). Periodic monitoring of P450 enzyme concentration by CO-reduced spectrum analysis was done to determine the optimum expression time of the different genes[2].

#### Purification of CYP6P4 and AgCPR from bacterial cultures

At the optimum expression time, 20-24 hours for CYP6P4, the cultures were stopped by incubation on ice for 10 minutes. They were transferred into centrifuge bottles and centrifuged at 2800g for 20 min at 4°C to pellet the cells. The supernatant was discarded and the pellet re-suspended in 20 ml ice-cold 1X TSE buffer (50mM Tris-acetate, 250mM sucrose and 0.25mM EDTA, pH 7.6). To digest the bacterial cell wall, 250 µl of 20mg/ml lysozyme was added to the re-suspended cells to a final concentration of 0.25 mg/mL and gently stirred at 4°C for one hour. The spheroplasts were pelleted at 2800g for 25 min at 4°C, the supernatant discarded and re-suspended in 8 ml ice-cold spheroplast resuspension buffer (100 mM potassium phosphate, 6 mM magnesium acetate, 20% (v/v) glycerol, and 0.1 mM dithiothreitol (DTT), pH 7.6. To the re-suspended spheroplasts, the following protease inhibitors were added; 10 mg/mL leupeptin, in water; 100 mM phenylmethylsulfonyl fluoride (PMSF), in isopropanol and 10 mg/mL of aprotinin, in 10 mM HEPES, pH 8.0 to a final concentration of 1 µg/mL, 1 mM and 1 µg/mL respectively. To lyse the spheroplasts membranes, sonication in 30 second bursts for a total of 2 minutes was carried out. The suspension was then centrifuged at 30, 000g for 20 minutes at 4°C to separate the membranes from other cell debris. The supernatant containing the membranes was transferred to ultracentrifuge tubes, topped up with buffer and centrifuged at 180, 000g for 1 hour at 4°C to pellet the membranes. The supernatant was discarded and 1ml of ice-cold 1X TSE buffer added to the membranes which were then dispersed using a Dounce tissue grinder (Wheaton). P450 concentration and AgCPR activity in the membrane was measured before being stored in -80°C.

#### Cytochrome P450 quantification

The concentration of active P450 was determined by comparing the spectra difference of reduced P450-carbon monoxide complex with reduced non-complexed P450. To monitor the concentration of P450 in cell cultures, 1 ml of culture was centrifuged at 16,400 rpm at 4°C to pellet the cells. The cells were gently re-suspended in 1 ml 1X P450 spectrum buffer (100 mM Tris-HCl, 10 mM CHAPS, 40% (v/v) glycerol, and 1 mM EDTA, pH 7.4) to avoid formation of bubbles and a few grains of sodium dithionate (reducing agent) added. The mixture was split into two spectrophotometer cuvettes and a baseline reading taken from 500 to 400 nm with a dual beam Cary 4000 UV-Vis spectrophotometer. The front cuvette was taken to a fume hood, bubbled with carbon monoxide at the rate of one bubble a minute for 50-80 seconds, mixed gently by inversion and replaced into the machine. The reference cuvette was also mixed by inversion before a second reading from 500 to 400 nm was taken. The concentration of active P450 was calculated by subtracting the peak measurements at 490 nm from that at 450 nm then dividing the difference with the extinction coefficient at 450 nm (0.091 µM/cm). In purified membranes, 50 µl of the membranes are added to 950 µl of 1X P450 spectrum buffer and quantified in a similar manner. However, the quantity of active P450 determined is multiplied by the dilution factor, in our case 20 to get the concentration of the enzyme per ml.

#### *Anopheles gambiae* P450 reductase quantification

We measured the quantity of AgCPR by estimating cytochrome c reductase (cyt c) activity. In a microtiter plate, 150 µl of 0.1 mM cyt C prepared in 0.3M potassium phosphate buffer at pH 7.7 was pipetted into six wells. Into each well, 2 µl of the membranes were added. In the first 3 wells, 150 µl of 0.3M potassium phosphate buffer was added to serve as control while 150 µl of 0.1 mM NADPH prepared using the same buffer was added in the other 3 wells to start the reaction. The reaction was monitored by taking readings at 550 nm every minute for 15 minutes with a microplate UV-Vis reader. The slope in mA550 per minute was calculated using a plot derived from OD measured per minute. To determine the specific activity of AgCPR (nmol of cyt c reduced/min/mg), the formula0.0552 * slope / protein concentration assuming a path-length of 0.85cm was used[2]. Protein concentration of the membranes was determined by Bradford assay with bovine serum albumin as the standard [3]. Diethoxyfluorescein (DEF) metabolism assay was also done as described previously to analyze P450 activity on the purified membranes[4].

#### In vitro insecticide metabolism assay

For each insecticide, 6 assays for the NADPH plus (test) reaction and 6 for the NADPH minus (negative control) were set up per biological replicate. This included 3 technical replicates for reaction with cytochrome b5 (b5) and 3 reactions without b5 in the test and control reactions. In an Eppendorf tube, 2 µl of 0.5 mM insecticide prepared in molecular grade absolute ethanol was added to 48 µl of enzyme mix containing the P450 enzyme with or without b5. To start the reaction, 50 µl of regeneration NADPH mix without NADP and that with NADP were added to the control and test Eppendorf tubes respectively. The tubes were incubated at 30°C on a shaking heating block at 1200 rpm for 2 hours. The reaction was quenched by adding 150 µl of HPLC grade acetonitrile. The tubes were inverted several times to ensure proper mixing of acetonitrile with the reaction mix and then centrifuged at 13,000 rpm for 20 minutes to pellet the membranes to prevent blockage of HPLC column. We transferred 150 µl of the supernatant into Thermofisher HPLC vials. HPLC run conditions were as previously described[5].Solvent ratio and monitoring wavelength were 85:15 Acetonitrile:water and 226nm for permethrin reactions and 80:20 Acetonitrile:water and 232nm for deltamethrin reactions.

### Appendix 3. Heterologous expression of *An. gambiae* CYP6AA1 and CPR using bac to bac expression system

#### Generation of Blunt-End PCR products

Full length coding sequences of *An. gambiae* CYP6AA1 Ugandan haplotype and *An. gambiae* CPR (transcript AY183375.1, [6] were amplified by PCR from the recombinant plasmid vectors pUAS:AgCYP6AA1 and pCW:AgCPR respectively, using the Phusion Hot Start II High-Fidelity DNA Polymerase (Thermo Scientific, Cat. No. F549S) and gene-specific primers (**Table 3.1**). Forward primers include a 5' minimal Kozak sequence (ACC), as in some cases this sequence acts as an enhancer, and reverse primers include the stop codon for expression of the native protein forms. Melting temperatures were calculated with Thermo's Multiple Primer Analyser software (https://www.thermofisher.com/uk/en/home/brands/thermo-scientific/molecular-biology/molecular-biology-learning-center/molecular-biology-resource-library/thermo-scientific-web-tools/tm-calculator.html) following polymerase manufacturer's instructions. PCR reactions contained 0.5uM primers, 200uM of each dNTP, 1U polymerase and 1uL plasmid DNA harbouring the template, in a final volume of 50uL. Cycling conditions for CYP6AA1 were: 98°C for 30 seconds followed by 25 cycles of 98°C for 10 seconds, 66°C for 20 seconds and 72°C for 30 seconds and a final extension step at 72°C for 10 minutes. Cycling conditions for AgCPR were: 98°C for 30 seconds followed by 25 cycles of 98°C for 10 seconds, 68°C for 30 seconds and 72°C for 30 seconds and a final extension step at 72°C for 15 minutes. PCR products were separated via gel electrophoresis and excised from gel using Qiagen's QIAquick Gel Extraction Kit (Cat. No. 28704). (**Figure 3.1**).

**Table 3.1. Primers used for the cloning of *An. gambiae* genes of interest**. In bold the start and end codons of transcription. ACC upstream the start codon is the minimal Kozak seq.

| Gene | Primer | Sequence (5'-3') | Prod. size (bp) |
|---|---|---|---|
| AgCYP6AA1 | AgUgCYP6AA1-F | ACC**ATG**GGTTACGTCAGTGTTGT | 1521 |
| (Ugandan Haplotype) | AgUgCYP6AA1-R | **TCA**CAGTTTCGTTGCGTTAAGCC | |
| AgCPR (transcript | AgCPR-F | ACC**ATG**GACGCCCAGACAGAAA | 2043 |
| AY183375.1), | AgCPR-R | **TTA**GCTCCACACGTCCGCCGA | |

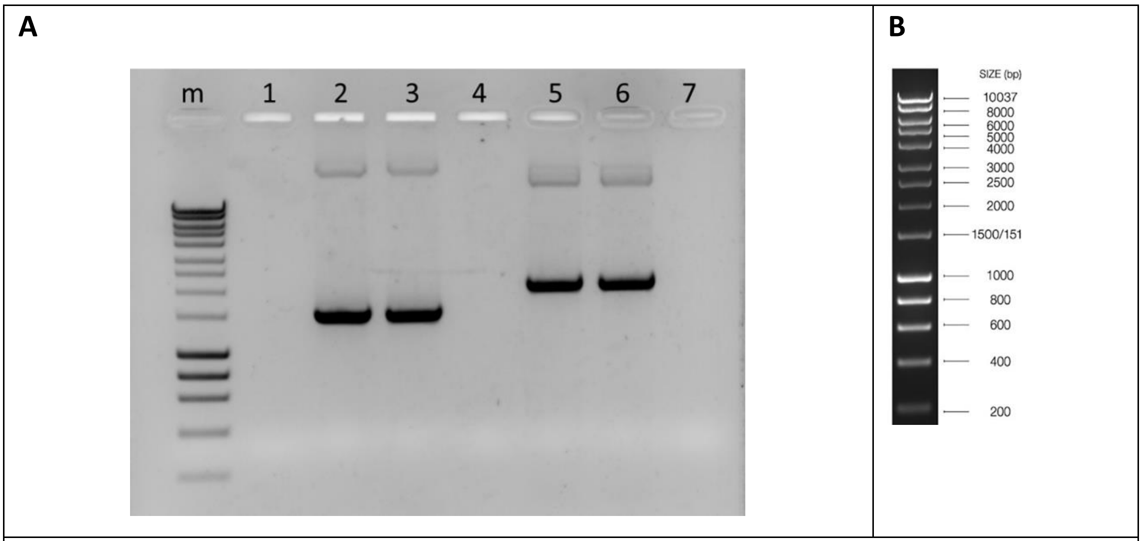

**Figure 3.1.** Molecular cloning of AgCYP6AA1 and AgCPR. **A.** Agarose gel results, PCR products after amplification with gene specific primers, m: HyperLadder 1kb-Bioline, 1:no sample, 2,3: AgCYP6AA1 (1521bp), 4:Non-template control, 5,6: AgCPR (2043bp), 7:Non-template control. **B** HyperLadder 1kb lanes.

#### Cloning to pFastBac/CT TOPO donor plasmid and analysis of positive clones

Purified Blunt-End PCR products were cloned into pFastBac/CT-TOPO vector (Invitrogen Cat.No. A11100), which contains a strong polyhedrin ($P_H$) promoter for high-level baculovirus-based protein expression in insect cells upstream the TOPO Cloning site and a mini-Tn7 elements for site-specific transposition into the bacmid DNA. Cloning reactions, consisting of 1ul of pFastBac/CT-TOPO vector plasmid vector, 1uL of freshly prepared purified PCR product, 1ul of Salt Solution in a final volume of 6uL, were incubated for 5 minutes at room temperature and then used to transform **One Shot Mach1-T1** competent cells. The resulting colonies were selected on ampicillin plates (100ug/mL) and positive clones were analyzed for correct orientation by colony PCR using one internal (gene specific forward primer) and one external primer (SV40 polyA Reverse primer). 10 colonies from each selective plate were analyzed (**Figure 3.2**). The PCR reactions contained 0.2uM primers, 200uM of each dNTP, 1U DreamTaq DNA polymerase and 1 colony as a template, in a final volume of 20uL. The cycling conditions for both CYP6AA1 and CPR were: 95°C for 3 minutes followed by 30 cycles of 95°C for 30 seconds, 42°C for 30 seconds and 72°C for 1 minute and a final extension step at 72°C for 10 minutes, with the annealing temperature calculated according to DreamTaq polymerase guide.

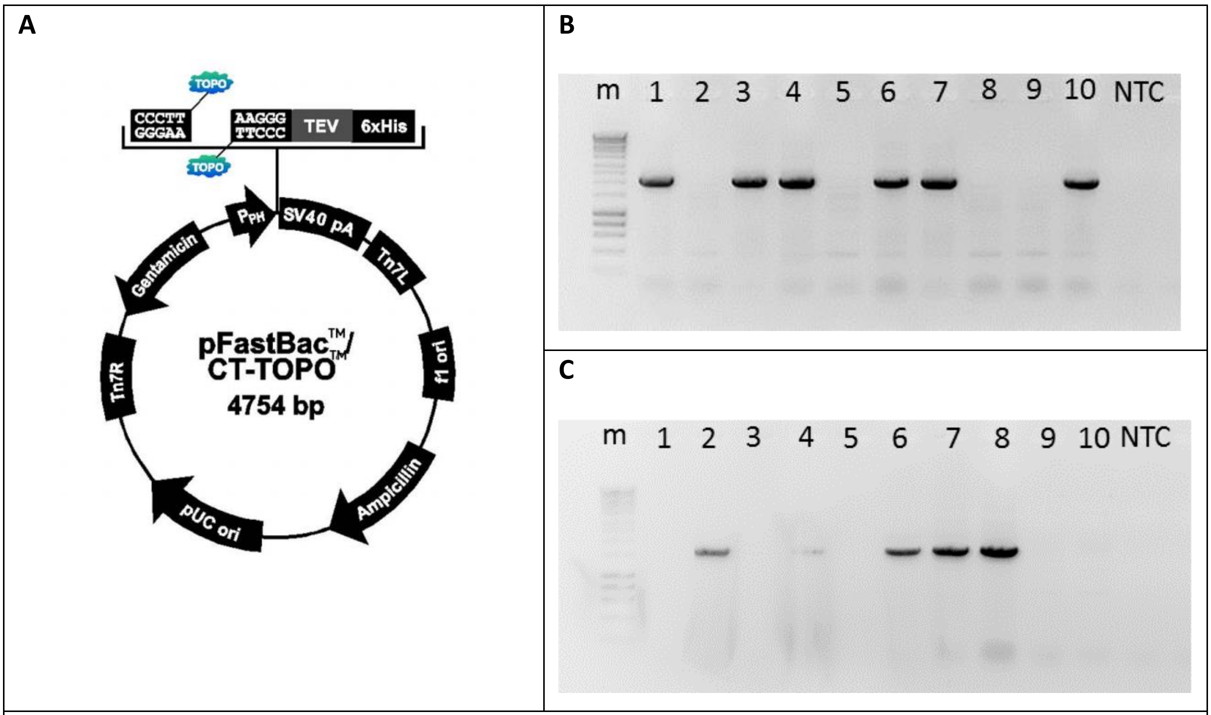

**Figure 3.2. A.** Map of the donor plasmid pFastBac/CT-TOPO, **B.** Colony PCR results for pFastBac:AgCPR using AgCPR-F and SV40 polyA R primer pair. Lanes 1, 3, 4, 6, 7, and 10 are positive clones. **C.** Colony PCR results for pFastBac:AgCYP6AA1 using AgUgCYP6AA1-F and SV40 polyA R primer pair. Lanes 2, 4, 6, 7 and 8 are positive clones. NTC: No-template control, m: HyperLadder 1kb.

Plasmids from positive clones (one per gene of interest) were grown in LB medium supplemented with 100ug/mL ampicillin, were extracted using QIAprep Spin Miniprep Kit, and were sent for sequencing to validate correct sequences (primers used for sequencing in **Table 3.2**). The sequences from the different reads were processed with BioEdit and were used to make the contigs containing the full sequences which were then aligned with the reference sequences. CYP6AA1 contig has full identity with the reference sequence (Alignment with the reference nucleotide sequence in **Figure 3.3**), whereas CPR contig has two SNPs in the coding sequence corresponding in two missense mutations (V230A and H459Y) in the protein sequence (Nucleotide and amino acid sequence alignment in **Figure 3.4** and 3.**5** respectively). However, these two SNPs correspond to real polymorphisms occurring in the population.

**Table 3.2. Primer pairs for the analysis of recombinant pFastBac plasmids and bacmids and internal gene specific primers used for sequencing.**

| **Application** | **Primer** | **Sequence (5'-3')** |
|---|---|---|
| analyzing recombinant plasmids | Polyhedrin Forward primer | AAATGATAACCATCTCGC |
| | SV40 polyA Reverse primer | GGTATGGCTGATTATGATC |
| analyzing recombinant bacmids | pUC/M13 Forward primer | CCCAGTCACGACGTTGTAAAACG |
| | pUC/M13 Reverse primer | AGCGGATAACAATTTCACACAGG |
| CPR internal primers | CPR 561 F | CGGCATCTACGTGGACAAGC |
| | CPR 1121 F | TCACCCACTATCTGGAGATAACG |
| | CPR 1680 R | TCACCCACTATCTGGAGATAACG |
| CYP6AA1 internal primer | AA1_qPCR_F1 | CCACGGTGAATCGGTTCGT |

```
CLUSTAL O(1.2.4) multiple sequence alignment

Contig-AA1col2            TAAAAAAACCTATAAATATTCCGGATTATTCATACCGTCCCACCATCGGGCGCGGATCCA        60
AA1_uganda_swept_hap      ------------------------------------------------------------        0

Contig-AA1col2            CCGGTCCCTTACCATGGGTTACGTCAGTGTTGTGTTATTTCTGGTGGTCCCCGCGCTCAC        120
AA1_uganda_swept_hap      -------------ATGGGTTACGTCAGTGTTGTGTTATTTCTGGTGGTCCCCGCGCTCAC        47
                                       ***********************************************

Contig-AA1col2            CCTGCTTTACATCTACTTGAAGCAGCACTACCGTCATTGGGCAAATCGGAACCTACCCCA        180
AA1_uganda_swept_hap      CCTGCTTTACATCTACTTGAAGCAGCACTACCGTCATTGGGCAAATCGGAACCTACCCCA        107
                          ************************************************************

Contig-AA1col2            GCTGGAAGCGTCCTTCCCGCTGGGCAACATGAAGGGCGTCGGGTCCGAGATTCACTTCAA        240
AA1_uganda_swept_hap      GCTGGAAGCGTCCTTCCCGCTGGGCAACATGAAGGGCGTCGGGTCCGAGATTCACTTCAA        167
                          ************************************************************

Contig-AA1col2            CGATGTGCTGAACGAAGCGTACGCAAAAGGTAAAGCACAGTCGGCCCCACTCGTCGGGCT        300
AA1_uganda_swept_hap      CGATGTGCTGAACGAAGCGTACGCAAAAGGTAAAGCACAGTCGGCCCCACTCGTCGGGCT        227
                          ************************************************************

Contig-AA1col2            GTACTTTATGCTCAAGCCCATACTGATCGTGACCGAGCTGGACATGGTGAAGCGCATACT        360
AA1_uganda_swept_hap      GTACTTTATGCTCAAGCCCATACTGATCGTGACCGAGCTGGACATGGTGAAGCGCATACT        287
                          ************************************************************

Contig-AA1col2            GGTGAAAGACTTTAGCAGCTTCCACGACCGTGGCCTGTACGTGAACGAGCGGGACGATCC        420
AA1_uganda_swept_hap      GGTGAAAGACTTTAGCAGCTTCCACGACCGTGGCCTGTACGTGAACGAGCGGGACGATCC        347
                          ************************************************************

Contig-AA1col2            CCTGTCCGGGCATCTGTTTGCGCTGGACGGTGAGCGGTGGCGCTATCTGCGCAACAAGCT        480
AA1_uganda_swept_hap      CCTGTCCGGGCATCTGTTTGCGCTGGACGGTGAGCGGTGGCGCTATCTGCGCAACAAGCT        407
                          ************************************************************

Contig-AA1col2            CAGCCCTACGTTCACGTCCGGCAAGATCAAGCTGATGTTCAGCACGATTTGCGAGATCGG        540
AA1_uganda_swept_hap      CAGCCCTACGTTCACGTCCGGCAAGATCAAGCTGATGTTCAGCACGATTTGCGAGATCGG        467
                          ************************************************************

Contig-AA1col2            TGACGAGTTTCTGGCCACGGTGAATCGGTTCGTCGAGCGGGATGAGCCGTTGGATGTGAA        600
AA1_uganda_swept_hap      TGACGAGTTTCTGGCCACGGTGAATCGGTTCGTCGAGCGGGATGAGCCGTTGGATGTGAA        527
                          ************************************************************

Contig-AA1col2            GCTGCTGTCACAGTGCTTCACCTGCGACGTGGTAGGCTCGTGCGCGTTCGGGCTGCAGTG        660
AA1_uganda_swept_hap      GCTGCTGTCACAGTGCTTCACCTGCGACGTGGTAGGCTCGTGCGCGTTCGGGCTGCAGTG        587
                          ************************************************************

Contig-AA1col2            CAACTCGCTCAAAAACGGAGGCTCCAAGCTGCTCGAGATCGGCGACAAGGTGTTCAAACC        720
AA1_uganda_swept_hap      CAACTCGCTCAAAAACGGAGGCTCCAAGCTGCTCGAGATCGGCGACAAGGTGTTCAAACC        647
                          ************************************************************

Contig-AA1col2            ACCTGCGTGGCGAAACATGCTCGCGTTTGCGCTCATTTCGTTTAAGAAGCTGGCGAAACG        780
AA1_uganda_swept_hap      ACCTGCGTGGCGAAACATGCTCGCGTTTGCGCTCATTTCGTTTAAGAAGCTGGCGAAACG        707
                          ************************************************************

Contig-AA1col2            ATTGCGCCTACCCGTGCTACCGGGCGAGGTGACGTCATTCTTTATGCCGCTTGTCACCGA        840
AA1_uganda_swept_hap      ATTGCGCCTACCCGTGCTACCGGGCGAGGTGACGTCATTCTTTATGCCGCTTGTCACCGA        767
                          ************************************************************

Contig-AA1col2            GACGGTGGCGGATCGGGAACGGAACTCGATCGAACGGCCCGATTTTCTCAATCTGCTGAT        900
AA1_uganda_swept_hap      GACGGTGGCGGATCGGGAACGGAACTCGATCGAACGGCCCGATTTTCTCAATCTGCTGAT        827
                          ************************************************************

Contig-AA1col2            ACAGCTCAAGAACAAAGGTACGGTCGAGGACGAATCAACGGAGGGACTGCAAAAGCTAAC        960
AA1_uganda_swept_hap      ACAGCTCAAGAACAAAGGTACGGTCGAGGACGAATCAACGGAGGGACTGCAAAAGCTAAC        887
                          ************************************************************

Contig-AA1col2            GCTCGACGAGGTGGCAGCGCAGGCGTTCGTGTTCTTCTTCGCCGGGTTCGAAACGTCCTC        1020
AA1_uganda_swept_hap      GCTCGACGAGGTGGCAGCGCAGGCGTTCGTGTTCTTCTTCGCCGGGTTCGAAACGTCCTC        947
                          ************************************************************

Contig-AA1col2            GACAACGCTCTCGTTCGCACTGTTCGAGTTGGCCAACAATCCCGACATTCAGGAGCGAGT        1080
AA1_uganda_swept_hap      GACAACGCTCTCGTTCGCACTGTTCGAGTTGGCCAACAATCCCGACATTCAGGAGCGAGT        1007
                          ************************************************************

Contig-AA1col2            GCGGGCGGAGGTGCTGGATAAGCTTCAACTCCACGACGGCAAGATAACGTACGACGCACT        1140
AA1_uganda_swept_hap      GCGGGCGGAGGTGCTGGATAAGCTTCAACTCCACGACGGCAAGATAACGTACGACGCACT        1067
```

```
                         ************************************************************
Contig-AA1col2           GAAGGAGATGATGTACCTCGATCAGGTCATCAATGAGACACTTCGTATGTATCCGCCAGT       1200
AA1_uganda_swept_hap     GAAGGAGATGATGTACCTCGATCAGGTCATCAATGAGACACTTCGTATGTATCCGCCAGT       1127
                         ************************************************************

Contig-AA1col2           TCCGCAGCTGATACGCGTCACCACTCAACCGTACAAGGTGGAGGGAGCGAACGTAACGCT       1260
AA1_uganda_swept_hap     TCCGCAGCTGATACGCGTCACCACTCAACCGTACAAGGTGGAGGGAGCGAACGTAACGCT       1187
                         ************************************************************

Contig-AA1col2           CGAGCCCGACACGATGCTGATGATACCGATCTACGCGATACATCACGATGCCAGCATCTA       1320
AA1_uganda_swept_hap     CGAGCCCGACACGATGCTGATGATACCGATCTACGCGATACATCACGATGCCAGCATCTA       1247
                         ************************************************************

Contig-AA1col2           TCCCGACCCGGAACGTTTCGATCCGGATCGGTTTGCACTGGCCGCGACGCACGCAAGGCA       1380
AA1_uganda_swept_hap     TCCCGACCCGGAACGTTTCGATCCGGATCGGTTTGCACTGGCCGCGACGCACGCAAGGCA       1307
                         ************************************************************

Contig-AA1col2           CACTCACGCGTTCCTGCCGTTCGGCGATGGGCCGCGGAATTGCATCGGCATGCGCTTTGG       1440
AA1_uganda_swept_hap     CACTCACGCGTTCCTGCCGTTCGGCGATGGGCCGCGGAATTGCATCGGCATGCGCTTTGG       1367
                         ************************************************************

Contig-AA1col2           GCTGCTGGAGGTAAAGTTTGGCATCGTGCAGATGGTGAGCAAGCTGCGGTTCACCGTCAG       1500
AA1_uganda_swept_hap     GCTGCTGGAGGTAAAGTTTGGCATCGTGCAGATGGTGAGCAAGCTGCGGTTCACCGTCAG       1427
                         ************************************************************

Contig-AA1col2           CGAGCGTATGCAGATGCCGCTCCGGATGTCGAAAACGGCCTCCATACTGGTGGCGGAAGG       1560
AA1_uganda_swept_hap     CGAGCGTATGCAGATGCCGCTCCGGATGTCGAAAACGGCCTCCATACTGGTGGCGGAAGG       1487
                         ************************************************************

Contig-AA1col2           GGGCATTTGGCTTAACGCAACGAAACTGTGAAAGGGCGAAAACTTGTACTTTCAAGGCCA       1620
AA1_uganda_swept_hap     GGGCATTTGGCTTAACGCAACGAAACTGTGA-----------------------------       1518
                         ******************************

Contig-AA1col2           TCACCATCACCATCACTAGCTCGAGGCATGC   1651
AA1_uganda_swept_hap     ------------------------------   1518
```

**Figure 3.3.** Nucleotide sequence from CYP6AA1 positive clone aligned with the reference sequence.

```
CLUSTAL O(1.2.4) multiple sequence alignment

Contig_AgCPR_col1      GTTTTGTAATAAAAAAACCTATAAATATTCCGGATTATTCATACCGTCCCACCATCGGGC   60
AY183375.1             ------------------------------------------------------------   0

Contig_AgCPR_col1      GCGGATCCACCGGTCCCTTACCATGGACGCCCAGACAGAAACGGAAGTGCCCGCGGGAAG   120
AY183375.1             ----------------------ATGGACGCCCAGACAGAAACGGAAGTGCCCGCGGGAAG   38
                                             **************************************
Contig_AgCPR_col1      CGTGAGCGACGAACCGTTCCTCGGCCCGCTTGACATCGTCCTGCTCGTCAGTCTGCTGGC   180
AY183375.1             CGTGAGCGACGAACCGTTCCTCGGCCCGCTTGACATCGTCCTGCTCGTCAGTCTGCTGGC   98
                       ************************************************************
Contig_AgCPR_col1      CGGCACTGCCTGGTATCTGCTCAAGGGCAAGAAAAAGGAAAGCCAAGCTAGTCAGTTTAA   240
AY183375.1             CGGCACTGCCTGGTATCTGCTCAAGGGCAAGAAAAAGGAAAGCCAAGCTAGTCAGTTTAA   158
                       ************************************************************
Contig_AgCPR_col1      ATCCTACTCGATCCAGCCGACGACGGTGAACACGATGACGATGGTGGAGAACTCGTTCAT   300
AY183375.1             ATCCTACTCGATCCAGCCGACGACGGTGAACACGATGACGATGGTGGAGAACTCGTTCAT   218
                       ************************************************************
Contig_AgCPR_col1      CAAGAAGCTACAGTCCTCGGGCCGCCGGCTCGTAGTGTTTTACGGTTCCCAAACAGGCAC   360
AY183375.1             CAAGAAGCTACAGTCCTCGGGCCGCCGGCTCGTAGTGTTTTACGGTTCCCAAACAGGCAC   278
                       ************************************************************
Contig_AgCPR_col1      GGCAGAGGAATTTGCCGGTCGCCTGGCGAAGGAAGGAATCCGCTACCAAATGAAGGGCAT   420
AY183375.1             GGCAGAGGAATTTGCCGGTCGCCTGGCGAAGGAAGGAATCCGCTACCAAATGAAGGGCAT   338
                       ************************************************************
Contig_AgCPR_col1      GGTCGCCGACCCAGAGGAGTGCAATATGGAAGAGCTGCTGATGCTGAAGGACATCGACAA   480
AY183375.1             GGTCGCCGACCCAGAGGAGTGCAATATGGAAGAGCTGCTGATGCTGAAGGACATCGACAA   398
                       ************************************************************
Contig_AgCPR_col1      ATCGTTGGCCGTGTTTTGCTTAGCGACGTACGGCGAGGGCGACCCGACGGACAACTGCAT   540
AY183375.1             ATCGTTGGCCGTGTTTTGCTTAGCGACGTACGGCGAGGGCGACCCGACGGACAACTGCAT   458
                       ************************************************************
Contig_AgCPR_col1      GGAGTTCTACGACTGGATTCAAAACAACGATCTAGATATGACCGGTTTGAATTACGCGGT   600
AY183375.1             GGAGTTCTACGACTGGATTCAAAACAACGATCTAGATATGACCGGTTTGAATTACGCGGT   518
                       ************************************************************
Contig_AgCPR_col1      GTTTGGCCTTGGCAACAAAACGTACGAGCACTACAACAAGGTCGGCATCTACGTGGACAA   660
AY183375.1             GTTTGGCCTTGGCAACAAAACGTACGAGCACTACAACAAGGTCGGCATCTACGTGGACAA   578
                       ************************************************************
Contig_AgCPR_col1      GCGTCTCGAGGAGCTCGGCGCGAACAGAGTGTTTGAGCTCGGTCTGGGTGACGATGATGC   720
AY183375.1             GCGTCTCGAGGAGCTCGGCGCGAACAGAGTGTTTGAGCTCGGTCTGGGTGACGATGATGC   638
                       ************************************************************
Contig_AgCPR_col1      CAACATCGAGGACTACTTCATCACGTGGAAGGAAAAGTTTTGGCCCACGGCTTGCGACTA   780
AY183375.1             CAACATCGAGGACTACTTCATCACGTGGAAGGAAAAGTTTTGGCCCACGGTTTGCGACTA   698
                       ************************************************** *********
Contig_AgCPR_col1      CTTCGGCATCGAGAGCACGGGCGAGGATGTGCTGATGCGGCAGTACCGCCTGCTGGAGCA   840
AY183375.1             CTTCGGCATCGAGAGCACGGGCGAGGATGTGCTGATGCGGCAGTACCGCCTGCTGGAGCA   758
                       ************************************************************
Contig_AgCPR_col1      GCCGGACGTGAGCGCGGACCGCATCTACACCGGCGAGGTGGCCCGGCTCCATTCGCTCCA   900
AY183375.1             GCCGGACGTGAGCGCGGACCGCATCTACACCGGCGAGGTGGCCCGGCTCCATTCGCTCCA   818
                       ************************************************************
Contig_AgCPR_col1      GACGCAGCGGCCACCGTTCGACGCGAAGAACCCGTTCCTCGCCCCGATCAAGGTGAACCG   960
AY183375.1             GACGCAGCGGCCACCGTTCGACGCGAAGAACCCGTTCCTCGCCCCGATCAAGGTGAACCG   878
                       ************************************************************
Contig_AgCPR_col1      GGAGCTGCACAAGGCGGGCGGCCGGTCCTGCATGCACGTCGAGTTCGACATCGAGGGCTC   1020
AY183375.1             GGAGCTGCACAAGGCGGGCGGCCGGTCCTGCATGCACGTCGAGTTCGACATCGAGGGCTC   938
                       ************************************************************
Contig_AgCPR_col1      GAAGATGCGGTACGAGGCGGGCGACCATCTCGCGATGTACCCGGTGAACGATCGCGATCT   1080
```

```
AY183375.1           GAAGATGCGGTACGAGGCGGGCGACCATCTCGCGATGTACCCGGTGAACGATCGCGATCT   998
                     ************************************************************
Contig_AgCPR_col1    GGTCGAGCGGCTCGGCCGGCTGTGCAATGCCGAGCTCGATACGGTCTTCTCGCTCATCAA   1140
AY183375.1           GGTCGAGCGGCTCGGCCGGCTGTGCAATGCCGAGCTCGATACGGTCTTCTCGCTCATCAA   1058
                     ************************************************************
Contig_AgCPR_col1    CACCGACACGGACAGCAGCAAGAAGCATCCGTTCCCGTGCCCCACCACCTACCGGACCGC   1200
AY183375.1           CACCGACACGGACAGCAGCAAGAAGCATCCGTTCCCGTGCCCCACCACCTACCGGACCGC   1118
                     ************************************************************
Contig_AgCPR_col1    GCTCACCCACTATCTGGAGATAACGGCGCTGCCGCGCACCCACATCCTGAAGGAGCTGGC   1260
AY183375.1           GCTCACCCACTATCTGGAGATAACGGCGCTGCCGCGCACCCACATCCTGAAGGAGCTGGC   1178
                     ************************************************************
Contig_AgCPR_col1    CGAGTACTGCGGCGAGGAGAAGGACAAGGAGTTCCTGCGCTTCATCTCGTCGACCGCGCC   1320
AY183375.1           CGAGTACTGCGGCGAGGAGAAGGACAAGGAGTTCCTGCGCTTCATCTCGTCGACCGCGCC   1238
                     ************************************************************
Contig_AgCPR_col1    CGACGGCAAGGCGAAGTACCAGGAGTGGGTGCAGGACAGCTGCCGCAACATCGTGCACGT   1380
AY183375.1           CGACGGCAAGGCGAAGTACCAGGAGTGGGTGCAGGACAGCTGCCGCAACATCGTGCACGT   1298
                     ************************************************************
Contig_AgCPR_col1    GCTCGAGGACATCCCGTCCTGCCATCCGCCGATCGATCACGTGTGCGAGCTGCTGCCCCG   1440
AY183375.1           GCTCGAGGACATCCCGTCCTGCCATCCGCCGATCGATCACGTGTGCGAGCTGCTGCCCCG   1358
                     ************************************************************
Contig_AgCPR_col1    GCTGCAGCCCCGCTACTACTCGATCTCCTCCTCGTCCAAGCTGCACCCGACGACGGTGCA   1500
AY183375.1           GCTGCAGCCCCGCTACCACTCGATCTCCTCCTCGTCCAAGCTGCACCCGACGACGGTGCA   1418
                     **************** *******************************************
Contig_AgCPR_col1    CGTGACCGCGGTGCTGGTGAAGTACGAGACGAAGACGGGCCGGCTGAACAAGGGCGTCGC   1560
AY183375.1           CGTGACCGCGGTGCTGGTGAAGTACGAGACGAAGACGGGCCGGCTGAACAAGGGCGTCGC   1478
                     ************************************************************
Contig_AgCPR_col1    GACGACCTTCCTCGCGGAGAAGCACCCGAACGATGGGGAGCCGGCACCGCGCGTACCAAT   1620
AY183375.1           GACGACCTTCCTCGCGGAGAAGCACCCGAACGATGGGGAGCCGGCACCGCGCGTACCAAT   1538
                     ************************************************************
Contig_AgCPR_col1    CTTCATCCGCAAGAGCCAGTTCCGGTTGCCGCCCAAGCCGGAAACGCCCGTGATCATGGT   1680
AY183375.1           CTTCATCCGCAAGAGCCAGTTCCGGTTGCCGCCCAAGCCGGAAACGCCCGTGATCATGGT   1598
                     ************************************************************
Contig_AgCPR_col1    GGGGCCCGGCACCGGGCTGGCACCGTTCCGGGGCTTCATCCAGGAGCGGGACCACTGCAA   1740
AY183375.1           GGGGCCCGGCACCGGGCTGGCACCGTTCCGGGGCTTCATCCAGGAGCGGGACCACTGCAA   1658
                     ************************************************************
Contig_AgCPR_col1    GCAGGAGGGCAAGGAGATTGGCCAGACGACGCTGTACTTTGGCTGCCGCAAGCGCTCCGA   1800
AY183375.1           GCAGGAGGGCAAGGAGATTGGCCAGACGACGCTGTACTTTGGCTGCCGCAAGCGCTCCGA   1718
                     ************************************************************
Contig_AgCPR_col1    GGACTACATCTACGAGGATGAACTGGAAGACTACTCCAAGCGCGGCATCATCAACCTTCG   1860
AY183375.1           GGACTACATCTACGAGGATGAACTGGAAGACTACTCCAAGCGCGGCATCATCAACCTTCG   1778
                     ************************************************************
Contig_AgCPR_col1    CGTTGCGTTCTCGCGCGACCAGGAGAAGAAGGTGTACGTGACGCACCTGCTCGAGCAGGA   1920
AY183375.1           CGTTGCGTTCTCGCGCGACCAGGAGAAGAAGGTGTACGTGACGCACCTGCTCGAGCAGGA   1838
                     ************************************************************
Contig_AgCPR_col1    CTCGGACCTCATATGGAGCGTGATCGGCGAAAACAAGGGACACTTTTACATCTGCGGTGA   1980
AY183375.1           CTCGGACCTCATATGGAGCGTGATCGGCGAAAACAAGGGACACTTTTACATCTGCGGTGA   1898
                     ************************************************************
Contig_AgCPR_col1    TGCGAAAAAATATGGCCACCGATGTGCGAAACATTCTGCTGAAGGTCATCCGCTCGAAGGG   2040
AY183375.1           TGCGAAAAAATATGGCCACCGATGTGCGAAACATTCTGCTGAAGGTCATCCGCTCGAAGGG   1958
                     ************************************************************
Contig_AgCPR_col1    TGGGCTCAGCGAAACCGAGGCCCAGCAGTACATCAAAAAGATGGAAGCCCAAAAACGATA   2100
AY183375.1           TGGGCTCAGCGAAACCGAGGCCCAGCAGTACATCAAAAAGATGGAAGCCCAAAAACGATA   2018
                     ************************************************************
Contig_AgCPR_col1    CTCGGCGGACGTGTGGAGCTAAAAGGGCGAAAACTTGTACTTTCAAGGCCATCACCATCA   2160
AY183375.1           CTCGGCGGACGTGTGGAGCTAA--------------------------------------   2040
                     **********************
Contig_AgCPR_col1    CCATCACTAGCTCGAGGCATGC       2182
AY183375.1           ----------------------       2040
```

**Figure 3.4.** Nucleotide sequence from CPR positive clone aligned with the reference sequence. SNPs are shown in red.

```
CLUSTAL O(1.2.4) multiple sequence alignment

Contig_AgCPR_col1    MDAQTETEVPAGSVSDEPFLGPLDIVLLVSLLAGTAWYLLKGKKKESQASQFKSYSIQPT   60
AY183375.1           MDAQTETEVPAGSVSDEPFLGPLDIVLLVSLLAGTAWYLLKGKKKESQASQFKSYSIQPT   60
                     ************************************************************

Contig_AgCPR_col1    TVNTMTMVENSFIKKLQSSGRRLVVFYGSQTGTAEEFAGRLAKEGIRYQMKGMVADPEEC   120
AY183375.1           TVNTMTMVENSFIKKLQSSGRRLVVFYGSQTGTAEEFAGRLAKEGIRYQMKGMVADPEEC   120
                     ************************************************************

Contig_AgCPR_col1    NMEELLMLKDIDKSLAVFCLATYGEGDPTDNCMEFYDWIQNNDLDMTGLNYAVFGLGNKT   180
AY183375.1           NMEELLMLKDIDKSLAVFCLATYGEGDPTDNCMEFYDWIQNNDLDMTGLNYAVFGLGNKT   180
                     ************************************************************

Contig_AgCPR_col1    YEHYNKVGIYVDKRLEELGANRVFELGLGDDDANIEDYFITWKEKFWPTACDYFGIESTG   240
AY183375.1           YEHYNKVGIYVDKRLEELGANRVFELGLGDDDANIEDYFITWKEKFWPTVCDYFGIESTG   240
                     *************************************************.**********

Contig_AgCPR_col1    EDVLMRQYRLLEQPDVSADRIYTGEVARLHSLQTQRPPFDAKNPFLAPIKVNRELHKAGG   300
AY183375.1           EDVLMRQYRLLEQPDVSADRIYTGEVARLHSLQTQRPPFDAKNPFLAPIKVNRELHKAGG   300
                     ************************************************************

Contig_AgCPR_col1    RSCMHVEFDIEGSKMRYEAGDHLAMYPVNDRDLVERLGRLCNAELDTVFSLINTDTDSSK   360
```

```
AY183375.1            RSCMHVEFDIEGSKMRYEAGDHLAMYPVNDRDLVERLGRLCNAELDTVFSLINTDTDSSK   360
                      ************************************************************

Contig_AgCPR_col1     KHPFPCPTTYRTALTHYLEITALPRTHILKELAEYCGEEKDKEFLRFISSTAPDGKAKYQ   420
AY183375.1            KHPFPCPTTYRTALTHYLEITALPRTHILKELAEYCGEEKDKEFLRFISSTAPDGKAKYQ   420
                      ************************************************************

Contig_AgCPR_col1     EWVQDSCRNIVHVLEDIPSCHPPIDHVCELLPRLQPRYYSISSSSKLHPTTVHVTAVLVK   480
AY183375.1            EWVQDSCRNIVHVLEDIPSCHPPIDHVCELLPRLQPRYHSISSSSKLHPTTVHVTAVLVK   480
                      **************************************:*********************

Contig_AgCPR_col1     YETKTGRLNKGVATTFLAEKHPNDGEPAPRVPIFIRKSQFRLPPKPETPVIMVGPGTGLA   540
AY183375.1            YETKTGRLNKGVATTFLAEKHPNDGEPAPRVPIFIRKSQFRLPPKPETPVIMVGPGTGLA   540
                      ************************************************************

Contig_AgCPR_col1     PFRGFIQERDHCKQEGKEIGQTTLYFGCRKRSEDYIYEDELEDYSKRGIINLRVAFSRDQ   600
AY183375.1            PFRGFIQERDHCKQEGKEIGQTTLYFGCRKRSEDYIYEDELEDYSKRGIINLRVAFSRDQ   600
                      ************************************************************

Contig_AgCPR_col1     EKKVYVTHLLEQDSDLIWSVIGENKGHFYICGDAKNMATDVRNILLKVIRSKGGLSETEA   660
AY183375.1            EKKVYVTHLLEQDSDLIWSVIGENKGHFYICGDAKNMATDVRNILLKVIRSKGGLSETEA   660
                      ************************************************************

Contig_AgCPR_col1     QQYIKKMEAQKRYSADVWS 679
AY183375.1            QQYIKKMEAQKRYSADVWS 679
                      *******************
```

**Figure 3.5.** Amino acid sequence from CPR positive clone aligned with the reference sequence. The two missense mutations are shown in red.

#### Generation of Recombinant Bacmids

##### Transformation of DH10Bac competent cells

pFastBac vectors harbouring the genes of interest, pFastBac:AgCYP6AA1 and pFastBac:AgCPR, were transformed into **Max Efficiency DH10Bac** *E.coli* competent cells containing the bacmid - viral DNA. Cells were left to recover for 4 hours and $10^{-1}$, $10^{-2}$ and $10^{-3}$ dilutions of the transformants were plated in selective plates containing 50 ug/mL kanamycin (Thermo, 11815-024), 10ug/mL tetracycline (Sigma, 87128), 7ug/mL gentamycin (Thermo, 15750-060) and 100ug/mL X-gal and 40ug/mL IPTG for blue-white selection. The plates were incubated for 48 hours at 37°C.

##### Analysis of positive clones with colony PCR

Positive white colonies, containing the recombinant bacmids were selected (the insertion disrupts the expression of LacZ peptide), re-streaked to selective plates and analyzed via colony PCR using an external (PUC/M13 F) and an internal primer (gene specific reverse). Successful transposition results in a PCR product longer than 4kb. Due to the big amplicon size, LongAmp Hot Start Taq (M0534S, NEB) was used, and annealing temperatures were calculated using NEBs Tm calculator. PCR reactions contained 0.4uM primers, 300uM of each dNTP, 2.5U DNA polymerase and 1 colony as a template, in a final volume of 25uL. Cycling conditions were: 94°C for 30 seconds followed by 25 cycles of 94°C for 20 seconds, 60°C for 30 seconds and 72°C for 3:30 minutes and a final extension step at 72°C for 10 minutes. PCR products were separated via gel electrophoresis (**Figure 3.6**).

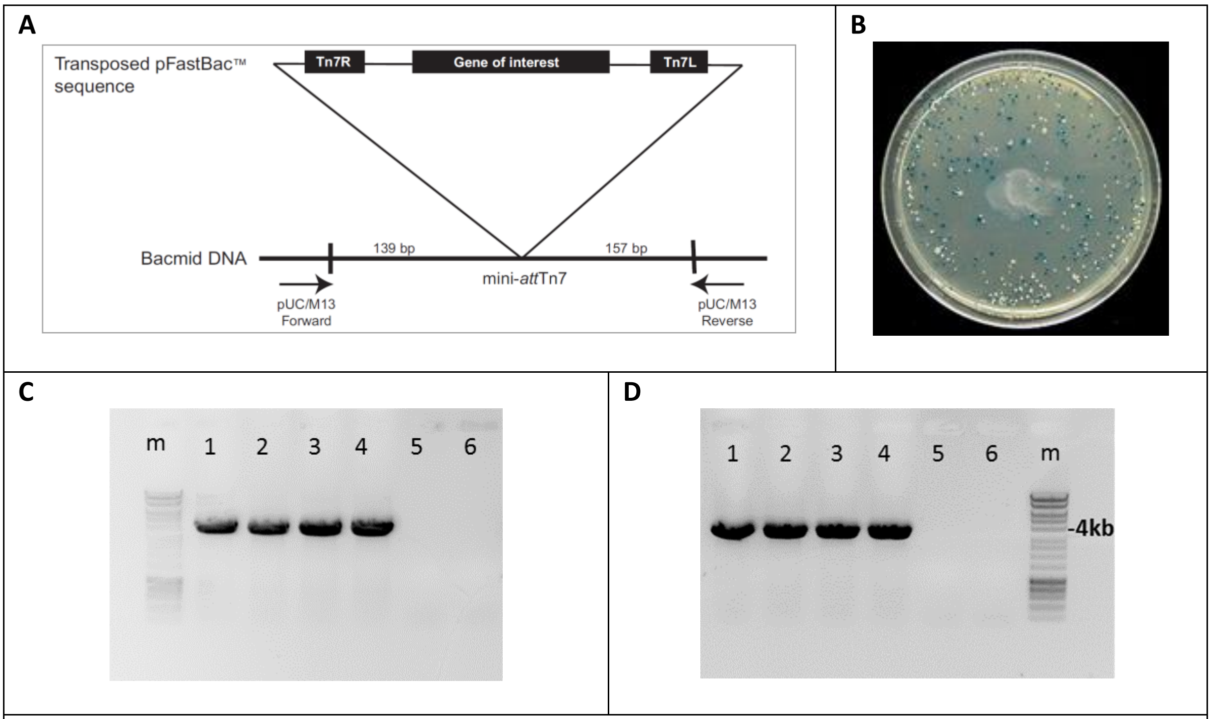

**Figure 3.6. A.** Mini-Tn7 elements for site-specific transposition into the bacmid DNA. **B.** Selective plate for blue/white selection. White colonies contain recombinant bacmids while blue colonies contain unaltered bacmid. **C.** Colony PCR results for AgCPR using pUC/M13 Forward and AgCPR-R primer pair. Lanes 1-4: positive clones, 5: No template control, 6: empty well. **D.** Colony PCR results for AgCYP6AA1 using pUC/M13 Forward and AgUgCYP6AA1-R primer pair. Lanes 1-4: positive clones, 5: No template control, 6: empty well. m: Bioline HyperLadder 1kb.

#### Isolation of Recombinant Bacmids

Positive colonies were grown in LB medium supplemented with 50 ug/mL kanamycin, 10ug/mL tetracycline and 7ug/mL gentamycin and recombinant bacmids were isolated using QIAprep Spin Miniprep Kit. Glycerol stocks of the recombinant bacmids in DH10Bac cultures were kept in -80°C. In order to obtain bacmid concentrations sufficient for transfection cultures of 200mL were grown and bacmids were isolated using PureLink HiPure Plasmid Maxiprep Kit (Thermo, K210006) and DNA was resuspended in 200ul TE. Bacmid concentrations are shown in the **Table 3.3** below. The resulting DNA was used to transfect *Sf*9 insect cells.

**Table 3.3. Bacmid isolation results**

| **r-bacmid** | **Concentration (ug/uL)** | **260/280** | **260/230** |
|---|---|---|---|
| AgCPR bacmid | 1.82 | 2.02 | 2.55 |
| AgCYP6AA1 bacmid | 1.59 | 2.03 | 2.55 |

#### Production of the recombinant baculovirus

##### Insect cell culture and transfection

*Spodoptera frugiperda Sf*9 cells were grown in Sf900 III SFM insect cell culture medium (Thermo, Cat.No 12658-019), in the absence of serum with 50ug/mL gentamycin (Thermo, Cat.No 15750-060) in adherent cultures, they were maintained at 27°C and sub-cultured weekly. Cells were established (cultured for 4 passages after thawing) and then they were transfected with the recombinant bacmids using ExpiFectamine Sf Transfection Reagent (Thermo, A38915).

Briefly, in a 6-well plate (Appleton Woods, BC010) 1 x $10^6$ cells were cultured per well in 3 mL medium. The next day, antibiotic containing Sf900 III SFM medium was removed and replaced by unsupplemented medium. For each transfection sample, 10uL of ExpiFectamine Sf Transfection Reagent were diluted in 250uL Opti-MEM I Reduced Serum Medium (Thermo, 31985062) and incubated for 5 minutes at room temperature. Subsequently 1ug bacmid DNA was added per sample. No DNA was added in the mock transfection control. See **Fig.3.7** for transfection plate layout. After 5 minutes incubation at room temperature the entire DNA-lipid mixture was added dropwise onto the cells and the plate was incubated at 27°C for 7 days. No signs of infection were obvious at that point.

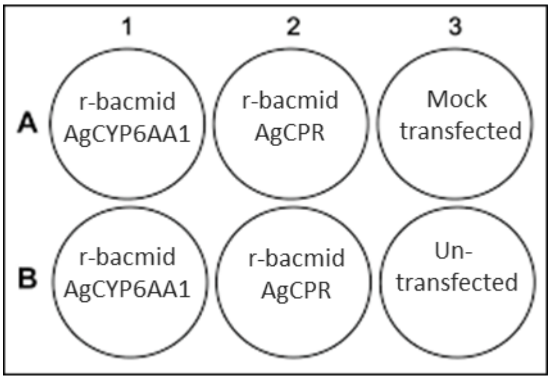

**Figure 3.7.** Transfection Layout in a 6-well plate.

##### Virus isolation and amplification

Seven days post transfection the medium containing the P0 viral stocks was collected by centrifugation (2000rpm, 5 minutes), filtered with 0.2um filters and used to infect Sf9 cells in two T25 flasks (Sigma, C6356-200EA) to amplify the viral stocks. One flask was infected with 200uL CYP6AA1-recombinant baculovirus (CYP6AA1rbv) and the other with 200uL CPR-recombinant baculovirus (CPRrbv), and they were inspected daily for visual signs of viral infection in a reverse phase microscope. Signs of successful viral infection include: increased cell diameter, cessation of cell, granular appearance, cell detachment and finally cell lysis. 8 days post infection, the medium containing the P1 viral stocks was collected by centrifugation (2000rpm, 5 minutes) and stored at 4°C. Infected cells were kept for a first expression test. Cells infected with CPRrbv will be checked for CPR expression using the cytochrome c assay and cells infected with CYP6AA1rbv can be assessed for P450 content and protein formation. In both assays the baseline will be estimated by untransfected cells as a control. P1 viral stocks will be used to identify the virus' multiplicity of infection (MOI) with plague assay, in order to progress with the expression experiments and validate the best ratio for CYP6AA1rbv and CPRrbv co-infection.

#### Metabolism assays

Deltamethrin and permethrin was dissolved in 100% ethyl alcohol to create 0.5 mM standard solution. The in vitro reactions contained 5 µM deltamethrin or permethrin, 100 pmol of CYP6AA1, 0.25mM $MgCl_2$, 0.1mM NADP+, 1mM glucose-6-phosphate and 1U/mL Glucose-6-phosphate dehydrogenase in 25mM potassium phosphate buffer (pH 7.4) in a total reaction volume of 200uL. NADP+ was not included in the control reactions. Reactions were performed in 5 replicates, and a two-sample T-test of sample reactions (+NADPH) vs the control (−NADPH) was performed for the statistical measurements of substrate depletion. After 2h incubation at 30 °C with 1200 rpm shaking, the reactions were stopped by adding 200µl of acetonitrile to the tubes, and then centrifuged for 20min at 13200xrpm to pellet the proteins. The supernatant was collected and transferred into HPLC glass vials for reverse-phase HPLC analysis. The mobile phase was 80% acetonitrile and 20% $H_2O$ deltamethrin and 85% acetonitrile and 15% $H_2O$ permethrin with flow rate: 1ml/min. The chromatographic analysis was conducted at 23°C and monitored by absorbance at 232nm deltamethrin and 226 nm permethrin.

### Appendix 4. Metabolism of pyrethroids by heterologously expressed P450s

#### Cyp6aa1

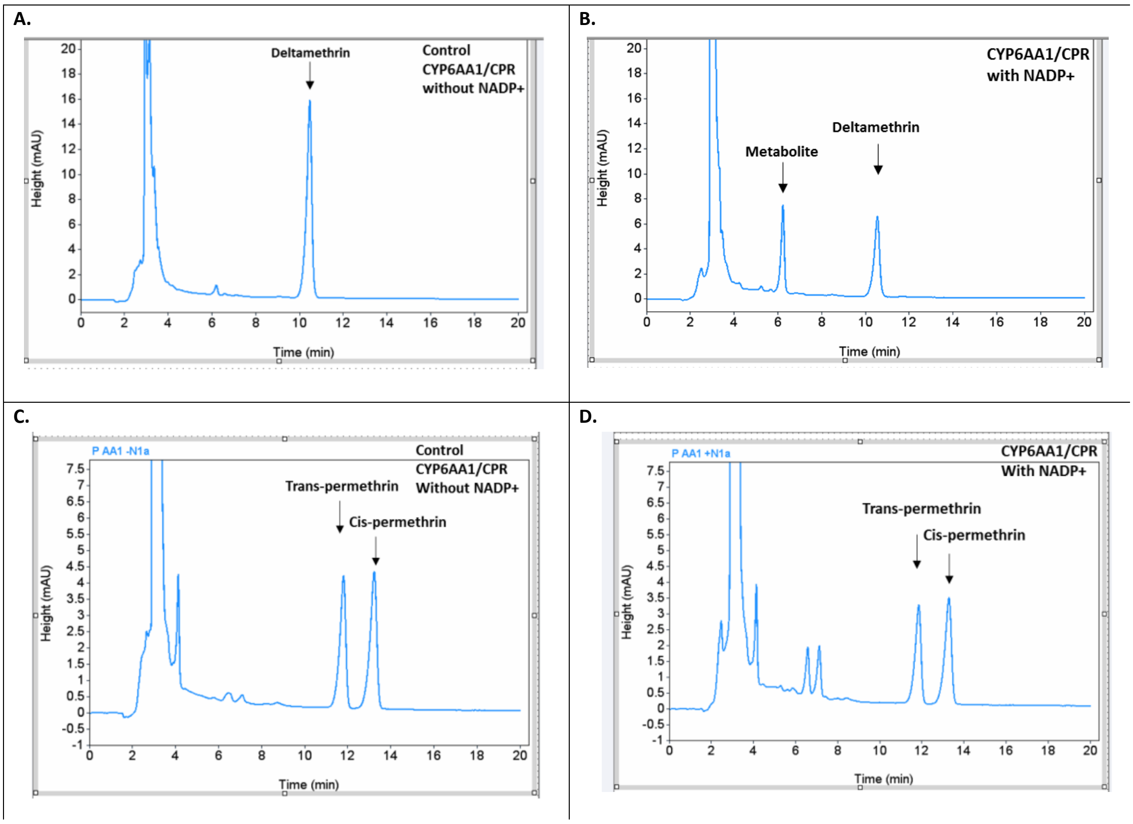

**Figure 4.1. Deltamethrin and permethrin metabolism by CYP6AA1/CPR cell pellets.** A. Deltamethrin (5 µM) incubated with CYP6AA1/CPR cell pellets for 2h at 30 °C without NADP+. B. Deltamethrin metabolized by Sf9 cell pellets expressing CYP6AA1/CPR with NADP+. Deltamethrin % depletion in the cell extracts expressing CYP6AA1/CPR with and without NADP+. (t-test: t= -9.67; d.f. = 8; P= 9.621E-06). C. Permethrin (5 µM) incubated with CYP6AA1/CPR cell pellets for 2h at 30 °C without NADP+. D.  Permethrin metabolized by Sf9 cell pellets expressing CYP6AA1/CPR with NADP+. The black arrows indicate the peaks for cis and trans permethrin D. Permethrin % depletion in the cell extracts expressing CYP6AA1/CPR with and without NADP+. ((t-test: t= -31.085; d.f. = 14; P= 2.554E-14)

### Appendix 5. *Cyp6aap* duplication frequency in collections from Cote d’Ivoire

| ***Cyp6aap* duplication**[1] | **Frequency 2012(n=71)** | **Frequency 2017 (n=164)**[2] | **P value**[3] |
|---|---|---|---|
| ***Cyp6aap-Dup7*** | 0.324 | 0.555 | 0.002 |
| ***Cyp6aap-Dup10*** | 0 | 0.05 | 1 |
| ***Cyp6aap-Dup11*** | 0.408 | 0.411 | 1 |
| ***Cyp6aap-Dup14*** | 0.465 | 0.742 | 0.00008 |
| ***Cyp6aap-Dup15*** | 0.394 | 0.331 | 0.4 |
| ***Any Cyp6aap-Dup*** | 0.972 | 0.976 | 1 |

**Table 5.1.** Cyp6aap duplication frequencies in specimens of Anopheles gambiae from Cote d’Ivoire collected in 2012 and 2017. [1]Duplication numbering as per [7]. [2] Sample sizes vary slightly for each Dup due to PCR failures [3]Fisher’s exact test

#### User Guides

- Bac-to-Bac™ TOPO™ Cloning Kit, Gibco
  - https://assets.thermofisher.com/TFS-Assets/LSG/manuals/bactobac_topo_cloning_kit_man.pdf
- Bac-to-Bac™ TOPO™ Expression System, Gibco
  - https://assets.thermofisher.com/TFS-Assets/LSG/manuals/MAN0000699_BactoBacTOPO_Expression_System_UG.pdf
- Growth and maintenance of insect cell lines, Gibco
  - http://tools.thermofisher.com/content/sfs/manuals/Insect_Cell_Lines_UG.pdf
- Guide to Baculovirus Expression Vector Systems (BEVS) and Insect Cell Culture Techniques
  - http://tools.thermofisher.com/content/sfs/manuals/bevtest.pdf
